# Dominant α-tubulin mutations rescue tauopathy neurodegenerative phenotypes in *C. elegans*

**DOI:** 10.64898/2026.03.18.712642

**Authors:** Sarah J. Benbow, Aleen D. Saxton, Misa Baum, Rikki L. Uhrich, Jade G. Stair, Kelly Keene, Chloe Dahleen, Linda Wordeman, Nicole F. Liachko, Rebecca L. Kow, Brian C. Kraemer

## Abstract

Tau protein, the primary component in neurofibrillary tangles characteristic of Alzheimer’s Disease and related dementia disorders, normally regulates microtubule growth and stability. While tau dysfunction contributes to the progression of tauopathies, the role of microtubules in disease has remained unclear. Through forward genetic screening in *Caenorhabditis elegans* tauopathy models, we found multiple tubulin gene mutations that rescue tau-mediated neurodegeneration. Whole animal behavioral and *in vitro* biochemical assays were employed to characterize mutation-driven effects on neuron function, neurodegeneration, and effects on tubulin and tau proteins as well as microtubule function. Mutant tubulin genes were found to confer different levels of suppression correlating with the level of mutant gene expression. Mutant tubulins did not drastically alter total tau protein levels, tau phosphorylation or aggregation, however tau-induced neurodegeneration was rescued. The suppression of tau toxicity by tubulin gene mutations cannot be explained by changes in tau or tubulin expression, tau phosphorylation, or tau aggregation state. Rather the tubulin mutations appear to act by influencing global microtubule properties. *In vitro* experiments using *C. elegans* tubulin in semi-isolated and isolated contexts have indicated changes to microtubule properties without observable changes to tau-tubulin affinity. This work suggests that manipulation of microtubules can rescue tauopathy even when pathological tau species persist, supporting the importance of understanding microtubule contributions to disease progression and investigation into microtubule targeted gene therapy or small molecule approaches for tauopathy intervention.

## Introduction

In dementia disorders that exhibit pathological tau protein, termed tauopathies, the microtubule associated protein tau becomes hyperphosphorylated, misfolded, and aggregates to form neurofibrillary and glial tangles [1, 2]. Tauopathy disorders include Down syndrome, corticobasal degeneration, progressive supranuclear palsy, Pick’s disease, Guam amyotrophic lateral sclerosis/Parkinson’s dementia complex, frontotemporal dementia with parkinsonism chromosome 17 type (FTDP-17) and Alzheimer’s disease[1, 2]. Autosomal dominant mutations in the MAPT gene encoding tau cause FTDP-17, indicating that abnormal tau accumulation can drive neuronal dysfunction and neurodegeneration independent of pathologies that may co-occur in other tauopathy disorders, and suggesting that abnormal tau may possess intrinsic toxicity [3–7]. The mechanistic underpinnings by which abnormal tau leads to neuronal dysfunction remain unclear. One hypothesis is that tau undergoes a gain of toxic function whereby the misfolding and oligomerization of tau creates toxic species [8–10]. A second hypothesis posits that as tau becomes hyperphosphorylated or misfolded, the normal microtubule binding activity is inhibited and results in loss of microtubule regulatory function [11, 12]. While mechanistically distinct, these hypotheses are not mutually exclusive, and both may contribute to a complex environment driving neuron dysfunction and decline.

Microtubules (MTs) are polymeric tubes composed of α- and β-tubulin subunit dimers that provide structural support to maintain the elongated structure of neurons as well as providing the tracks on which motor proteins move cargo between the cell body and the ends of axons and dendrites. MTs also undergo numerous post translational modifications and interactions with microtubule associated proteins (MAPs) to maintain balance between tightly controlled dynamic and stable phases [11, 13]. Multiple lines of evidence implicate MT dysfunction, manifesting as dendritic arbor simplification and disruption of axonal transport, as a component in neurodegenerative diseases [11, 13]. Similar to mammals, the *C. elegans* tubulin super-family of genes consist of multiple α and β isoforms. *C. elegans* have 9 α-tubulin genes and 6 β-tubulin genes expressed in tissue specific combinatorial ratios, providing precise regulatory control of a cell’s tubulin pool and resulting microtubule properties [14]. This differential expression is particularly notable in specialized cells like neurons which are highly polarized in their structure and function [15].

Tau plays a critical role in the regulation of the neuronal microtubule cytoskeleton. Tau binds directly to the microtubule lattice, conferring stability and at the plus tip, where it mediates the addition of new tubulin dimers particularly during microtubule rescue [16–19]. Tau’s affinity for MTs is regulated in part by the expression of 6 alternatively spliced isoforms varying in the number of N-terminal and microtubule binding domain repeats [20, 21]. Additionally, tau contains numerous serine, threonine and tyrosine residues which can be reversibly phosphorylated. The addition of phospho-groups at these sites disrupts tau-MT electrostatic interactions causing Tau to dissociate from the microtubule surface [12, 22–25]. Treatment of mice with Epothilone D (EpoD), a MT stabilizing drug, have prevented the development of tau pathology in young mice, cleared existing tau pathology in aged mice and improved cognitive performance [26, 27]. Additionally, EpoD treatment increased MT stability, density, increased fast axonal transport and decreased neuronal dystrophy in PS19 mice [26]. Recently another microtubule stabilizing compound, CNDR51657 was shown to reduce AT8 staining that detects disease-relevant tau phosphorylated at ser202 and ser205 and accumulation of TAR DNA-binding protein 43 (TDP-43) to ameliorate microtubule disruption in the optic tracts of repeated traumatic brain injury (rTBI) treated mice. Additionally, CNDR51657 improved memory and cognitive function in rTBI-treated mice [28]. Taken together, this evidence suggests increasing microtubule stability shows promise as a therapeutic strategy. However, these studies have not directly addressed the molecular underpinnings driving the neuronal protection seen with these compounds. *Caenorhabditis elegans* offers a number of key advantages as an experimental model for human disease namely, genetic homology to humans, rapid generation time, short lifespans, and tractable genetics. Expression of human tau in *C. elegans* neurons results in toxic phenotypes similar to those seen in human tauopathies including behavioral abnormalities, axonal degeneration, neuron loss, premature death and accumulation of phosphorylated and aggregated tau protein, which worsen with age [29]. Previously, we have employed these models in forward genetic screens to uncover genetic suppressors of tau pathology [30–36]. Using similar methodology in this study has revealed genetic mutations in 3 of the 9 *C. elegans* α-tubulin genes, *tba-1, tba-2,* and *mec-12*, that suppress pathological tau toxicity in our human-tau transgenic *C. elegans* models. We used *C. elegans* models to characterize mutant tubulin rescue of tauopathy phenotypes and probe the underlying molecular mechanisms. Tau phenotype suppressing mutations do not alter overall accumulation of pathological tau or alter overall tubulin expression. Rather, our results suggest mutations act by altering the properties of microtubules. *In vitro* experiments using semi-isolated and isolated tubulin protein indicate changes to microtubule properties without changing tau-tubulin binding affinity. The studies presented here demonstrate that a single amino acid change in tubulin can have profound effects on the pathological progression of tauopathy related neurodegeneration.

## Results

### Tau-phenotype suppressing mutations cluster in the tau binding region of α-tubulin

We previously generated a *C*. *elegans* model of tauopathy where a genomically integrated multicopy transgene array encoding human tau isoform with all 4 possible microtubule binding repeats, and 1 of 2 possible N-terminal repeats (4R1N) under the pan-neuronal *aex-3* promotor was integrated into the *C. elegans* genome [29]. In this model, human 4R1N tau protein accumulates in a highly phosphorylated state within neurons leading to tau aggregation, neuronal dysfunction manifesting as behavioral deficits, neurodegeneration, and decreased lifespan. To dissect molecular genetic pathways contributing to tauopathy phenotypes, we have employed a classical genetic mutagenesis and screening approach in a tau transgenic *C. elegans* searching for mutants resistant to tau mediated neurodegenerative phenotypes. Using this approach, we have previously isolated strong *<u>su</u>*ppressor of *t*auopathy (*sut*) mutations in the *dib-1, sut-1, sut-2, sut-6,* and *spop-1* genes [30–36].

Using the same screening paradigm, we uncovered several distinct genetic mutations in α-tubulin genes that rescue tau-induced motility deficits in *C. elegans* models of tauopathy. Eleven tau suppressing missense mutations have been identified in multiple tubulin genes. The isolated mutations cluster in the same region across three α-tubulin genes (*tba-1*, *tba-2*, *mec-12*) and are detailed in **Supplementary Table 1**. To date we have measured suppressive capability of several mutant strains: *mec-12* (D431N), *tba-1* (E437K), *tba-2* (D429A), *tba-2* (D429N), and *tba-2* (V433D). We selected *mec-12* (D431N), *tba-1* (E437K), and *tba-2* (D429N) for more extensive exploration as they encompassed the range of suppression observed, and further focused on the strongest suppressor: *tba-2* (D429N). Mutations in *mec-12, tba-1,* and *tba-2* occur at the C-terminal encoding region corresponding to helix 12 in each of these α-tubulin encoding genes (**Figure 1**). An alignment of the protein sequences (**Figure 1A**) revealed near identical homology in primary sequences across the various α-tubulin genes in *C. elegans* as well as near human homologues, indicating that the mutations occur in a highly conserved region of the proteins. Interestingly, helix 12 is exposed on the outside surface of MTs (depicted in **Figure 1B and 1C**) and is predicted to play a prominent role facilitating tau-MT binding [16], leading us to hypothesize that the mutations may confer suppression of tau-induced phenotypes through a mechanism of altered tau-tubulin and/or tau-MT binding affinity.

**Figure 1.**
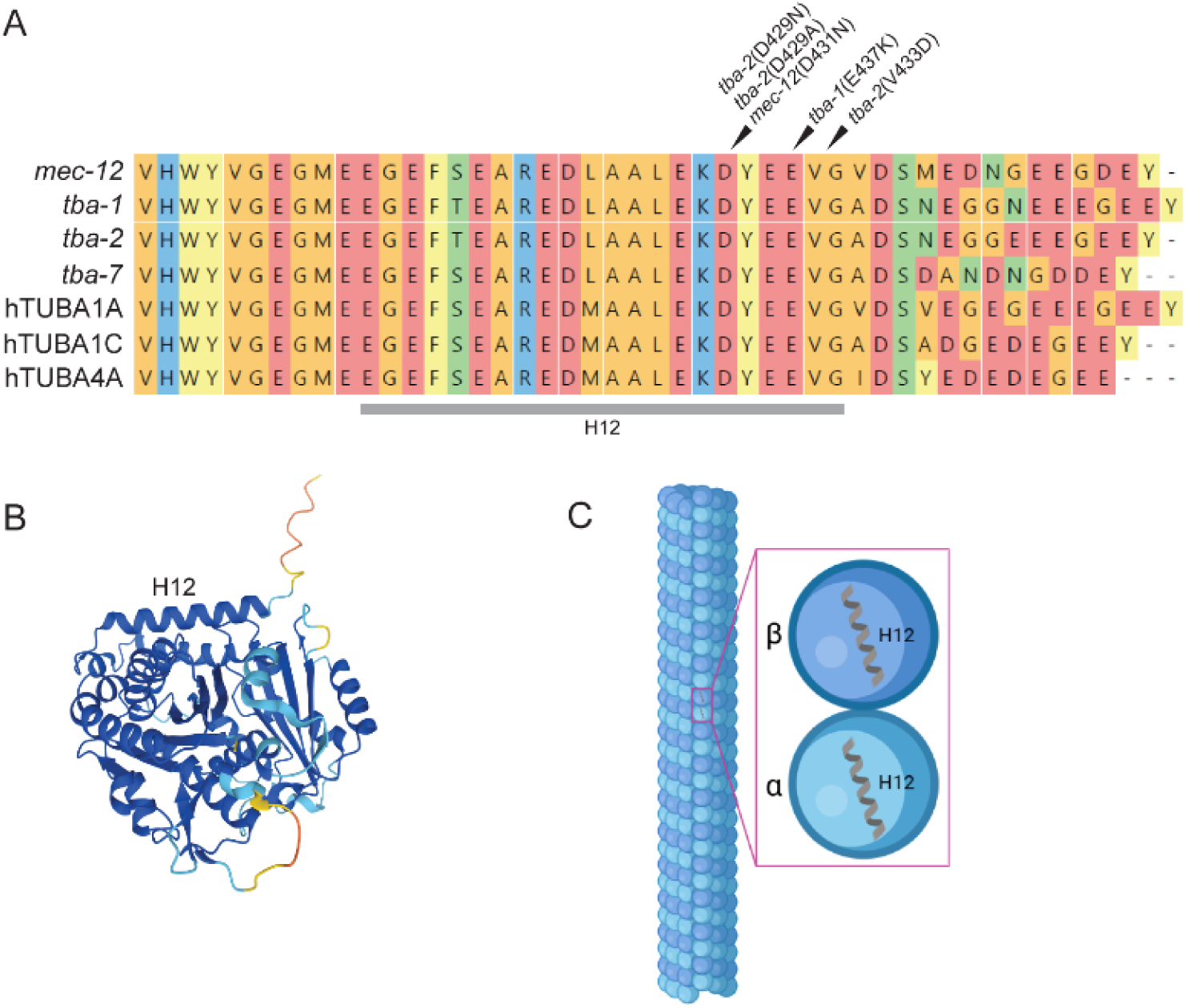
α-tubulin mutations cluster in conserved functionally important region on exterior microtubule surface. Alignment (A) of *C. elegans* alpha-tubulin genes *mec-12, tba-1, tba-2, tba-7* and human alpha tubulin homologues TUBA1A, TUBA1C, and TUBA4A demonstrate the high degree of conservation at the region corresponding to helix 12 seen in the AlphaFold2 predicted structure for *tba-2* UniProt Accession: P34690 (B) which are exposed to the outside surface of the MT when assembled and are involved in MAP (e.g. tau) binding (C) Created in BioRender. Wheeler, J. (2026) https://BioRender.com/ol82yg3.

### *C. elegans* α-tubulin mutations rescue tau-mediated neuronal dysfunction

To investigate the impact of mutations in α-tubulin genes on tau induced neuronal dysfunctions, we measured *C. elegans* behavioral deficits caused by transgenic tau expression in a swimming motility assay in which *C. elegans* are placed in buffer and motility is measured (body bends over time) as a readout of neuronal dysfunction. Human tau transgenic worms that express human tau pan-neuronally exhibit an uncoordinated locomotion phenotype driven by neuronal dysfunction that worsens with age, whereas tau-transgenic animals containing a tubulin mutation exhibit rescue of this neuronal dysfunction. We tested representatives from each mutant tubulin gene and observed that two mutant alleles of *tba-2* conferred the strongest levels of tau-induced behavioral deficit suppression, rescuing tau transgenic animals to wild type levels of motility. We observed that two of the mutations found in *tba-2* (D429N, and D429A) conferred the strongest levels of tau-induced motility deficit suppression, rescuing to wildtype or near wildtype levels (94% and 100% of wildtype levels respectively) of motility in a *C. elegans* model of tauopathy expressing high levels of human tau (WT-TauH, strain CK144) (**Figure 2 A-B**). Interestingly, mutant *mec-12* demonstrated the lowest level of suppression observed for the tubulin genes tested, conferring suppression to approximately 37% of wildtype levels (**Figure 2C**). We observed similar patterns of α-tubulin mediated suppression in a transgenic tauopathy model expressing more moderate levels of human tau (strain CK1443, WT-TauM, **Supplementary Figure 1**), with *tba-2* (V433D) mutations suppressing motility deficits in WT-TauM expressing worms to approximately 82% of wildtype levels. Additionally, we tested whether mutant *tba-2* could suppress the toxic effects conferred by the FTLD-tau (V337M) mutation (strain CK10, **Figure 2D**) [29]. Here, the same *tba-2* mutant provided only partial (∼87% of wildtype levels) rescue of deficits in the mutant tau-transgenic background (**Figure 2D**). The suppressor mutant found in *tba-1* also confers partial rescue (∼79% of wildtype levels) of neuronal dysfunction as measured in the motility assay (**Supplementary Figure 2**). The failure of an α-tubulin truncation (Q230Stop) mutant to suppress tau toxicity suggests that this suppression is specific to the amino acid substitutions within helix 12 (**Supplementary Figure 3**). Notably, when testing the level of suppression seen in *C. elegans* heterozygous for the *tba-2* (D429N)/+ mutation, we observed heterozygous *tba-2* D429N mutation rescues tauopathy phenotypes to approximately 65% of wildtype levels, demonstrating that mutant *tba-2* appears to be a strong semi-dominant suppressor of tauopathy neuronal dysfunction (**Supplementary Figure 4**).

**Figure 2.**
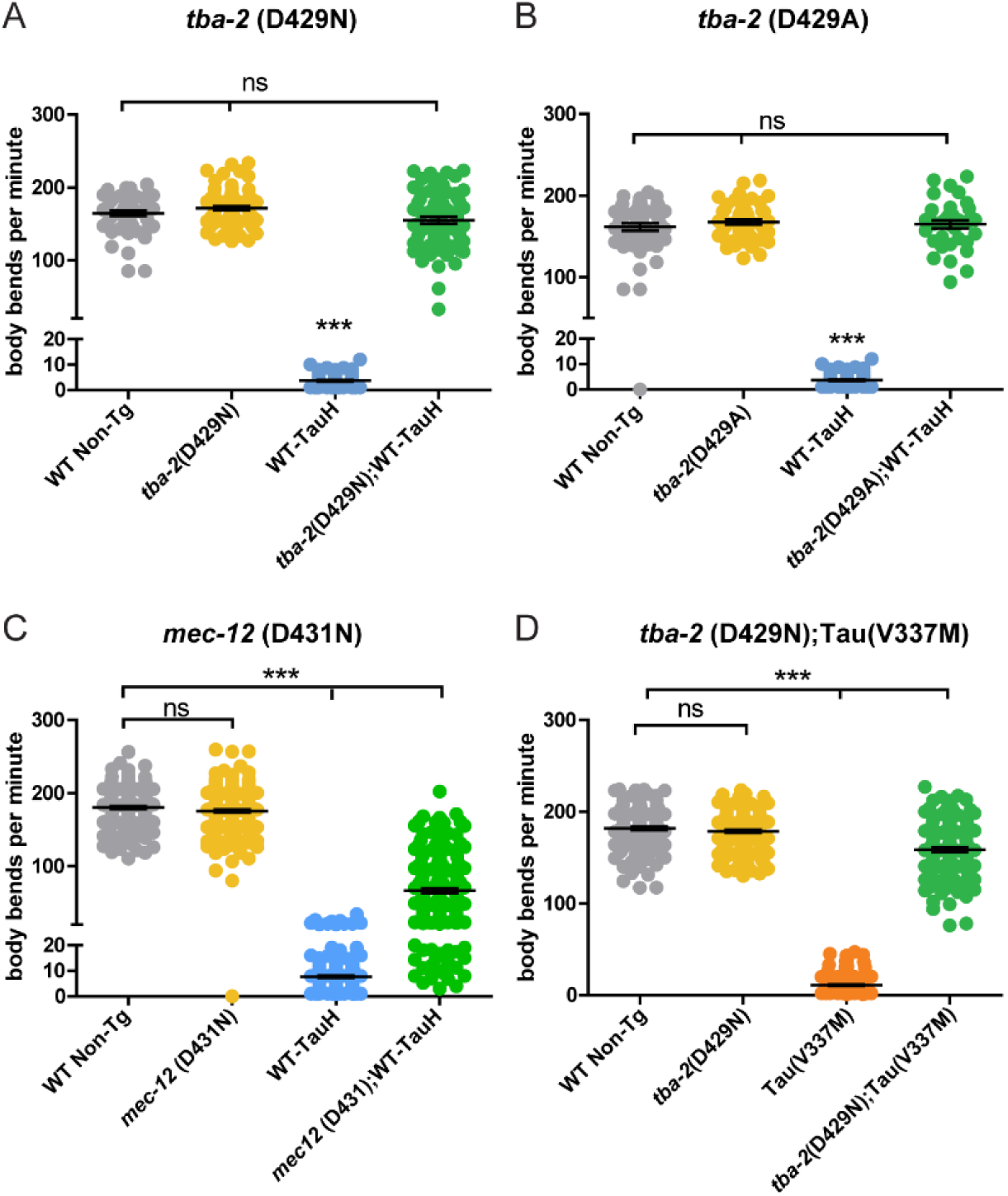
Mutations in *C. elegans* alpha-tubulin genes suppress wildtype tau-induced motility deficits to varying degrees. Independent *tba-2* mutant alleles (D429N and D429A) in high expressing wildtype human tau background (WT-TauH, strain CK144) rescue tau-induced motility deficits to wildtype levels (A and B, respectively), whereas only partial rescue is seen with similar mutations in *mec-12* (D431N) (C) (\*\*\**p<0.001,* error bars reflect SEM,*). tba-2* (D429N) also rescues mutant tau, Mut-TauH (strain CK10) induced deficits to ∼87% of wildtype levels (D, \*\*\**p<0.0001,* error bars reflect SEM). (A-B: n ≥ 55, C-D: n ≥142)

### Abundance of mutant α-tubulin drives the strength of tauopathy suppression

The well supported multi-tubulin hypothesis asserts that α- and β-tubulin isotypes are functionally distinct and differential spatial-temporal expression impart specific microtubule properties across different tissue types [37–40]. According to the *C. elegans* Neuronal Gene Expression Network (CeNGEN) dataset [41], *tba-*2 is the more abundantly expressed α-tubulin isotype in neurons as compared with *tba-1* and *mec-12*, with *mec-12* being the lowest expressed of these three genes neuronally. Simplistically, the degree of tubulin expression appears to align with the strength of tauopathy behavioral suppression as observed in the swimming assays. Thus, we hypothesized that the relative strength of tau-neurotoxicity suppression may be imparted by the relative levels of mutant α-tubulin isotype expression. To directly test this hypothesis, we generated integrated multicopy D429N mutant *tba-2* transgenic strains of *C. elegans* in which mutant *tba-2* transgenes are restricted to pan-neuronal expression. We selected mutant *tba-2* transgenic lines (*tba-2-D429N* Tg-A and *tba-2-D429N* Tg-B) that exhibited different levels of tau suppression when crossed into a tau-transgenic line expressing high levels of human tau (WT Tau-H, CK144) as assessed by motility deficits observed in the swimming assay described previously. We observed that the *tba-2-D429N* Tg-B animals exhibited significantly greater levels of tau induced motility deficit suppression (**Figure 3A**). Transgene expression analysis by qRT-PCR revealed transgenic expression of mutant *tba-2* was five-fold greater in *tba-2-D429N* Tg-B animals demonstrating that greater levels of mutant tubulin gene expression indeed correlated with a greater degree of motility deficit suppression (**Figure 3B**). As a control, we measured total, both mutant and endogenous, *tba-2* mRNA throughout the worm, which was not different between the two strains (**Supplementary Figure 5**).

**Figure 3.**
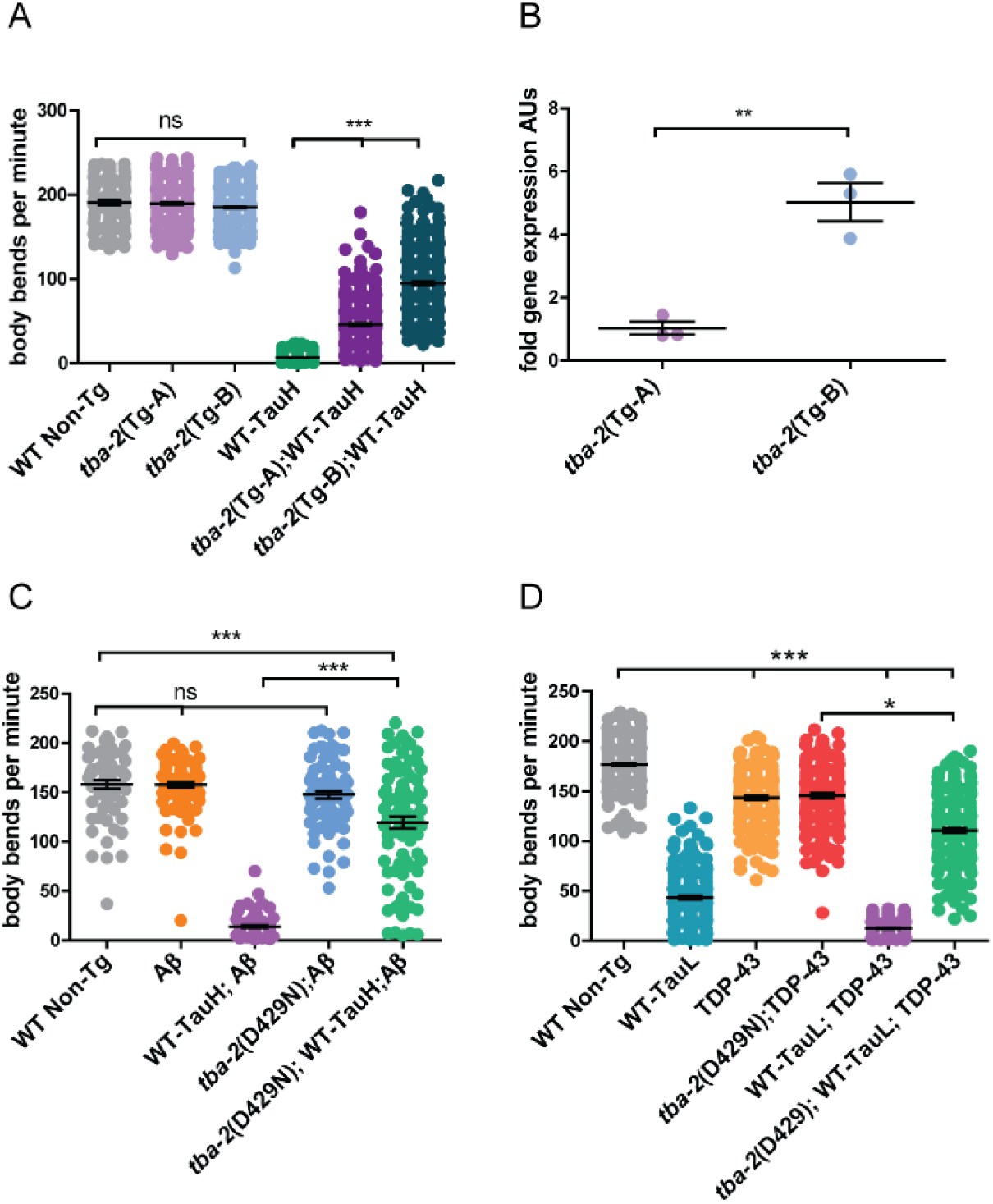
Tubulin expression correlates positively with expression level and suppresses in co-pathology models. Transgenic *C. elegans* with pan-neuronal over-expressing mutant *tba-2* (D429N) were created and compared for level of tau-induced motility deficit suppression. The strain *tba-2*(Tg-A) showed a lower level of motility deficit suppression as compared to *tba-2*(Tg-B) (A). qRT-PCR analysis revealed that *tba-2*(Tg-A) expressed a significantly lower amount of mutant tubulin as compared to *tba-2*(Tg-B) (B), correlating with the relative difference between the strain in motility rescue. Motility means were compared using a Kruskal-Wallis test with a Dunn’s post-hoc analysis, ***p<0.001, **p<0.01, n ≥ 110, error bars reflect SEM. qPCR data was analyzed using a Student’s T-test p=0.003, n=3, error bars represent SEM. Mutant tubulin partially suppresses motility deficits in worms with co-pathology models. *tba-2* (D429N) provides partial suppression of synergistic tau and Aβ pathology in worms expressing both human tau at high levels and toxic human Aβ peptide (C, \*\*\**p<0.001*, n ≥ 85, error bars reflect SEM). *tba-2* partially suppresses synergistic tau and TDP-43 pathology in worms expressing both wildtype human tau at lower levels and human TDP-43 protein (D, \**p<0.05,* \*\*\**p<0.0001* in a Kruskal-Wallis test with a Dunn’s post-hoc analysis, n ≥ 184 error bars reflect SEM).

### *tba-2* confers strong suppression of tau-Aβ and tau-TDP-43 co-pathology models

Tau neurofibrillary tangles and (Amyloid-β) Aβ plaques define Alzheimer’s disease in humans and act synergistically to enhance pathological phenotypes in animal models including tau/amyloid bigenic *C. elegans* [42]. Recent neuropathological data from human tissue has revealed that a number of AD cases additionally harbor aggregates of TDP-43 [43]. AD cases with TDP-43 co-pathology exhibit heterogeneous arrays of protein deposits and differ in the severity of toxic phenotypes. Tau and TDP-43 bigenic expression has also been shown to drive synergistic neuronal dysfunction and neurodegeneration in *C. elegans* [43]. Here we used animal co-pathology models to test the effects of *tba-2* suppression when tau phenotypes are enhanced by either Aβ peptide or TDP-43 protein expression. In both co-pathology models, mutant *tba-2* only strongly suppresses motility deficits to levels nearing the relative contribution of tau toxicity to the overall phenotype, ∼75% of wildtype motility in Aβ co-pathology model and ∼71% of wildtype motility (**Fig. 3C and 3D**).

### Mutant tubulin eliminates tau-mediated neurodegeneration

Transgenic *C. elegans* neuronally expressing high levels of human tau exhibit neurodegeneration that worsens with age [29]. We investigated whether *tba-2* and *mec-12* suppressed tau-induced neuronal loss by measuring the loss of GFP-marked GABAergic neurons in the ventral nerve cord [29, 44]. *C. elegans* expressing the human WT-Tau transgene along with mutant α-tubulin were compared to wildtype, *tba-2*, and tau-transgenic controls. Wildtype *C. elegans* develop the same number of neurons in each animal, facilitating the quantification of GFP marked neurons lost to neurodegeneration in mutant and transgenic strains. We observed the expected significant mean loss of ∼1.5 neurons by day 1 adulthood in WT-Tau worms (**Figure 4A, 4B**), whereas both *tba-2*;WT-Tau **(Figure 4A, 4C)** and *mec-12;*WT-Tau (**Figure 4B)** animals exhibited no significant neuron loss as compared to wildtype animals. These data suggest that both mutant *tba-2* and *mec-12* suppress tau-induced neuron loss and the differences observed in motility deficits result from differences in rescue of neuronal dysfunction rather than rescue of neurodegeneration.

**Figure 4.**
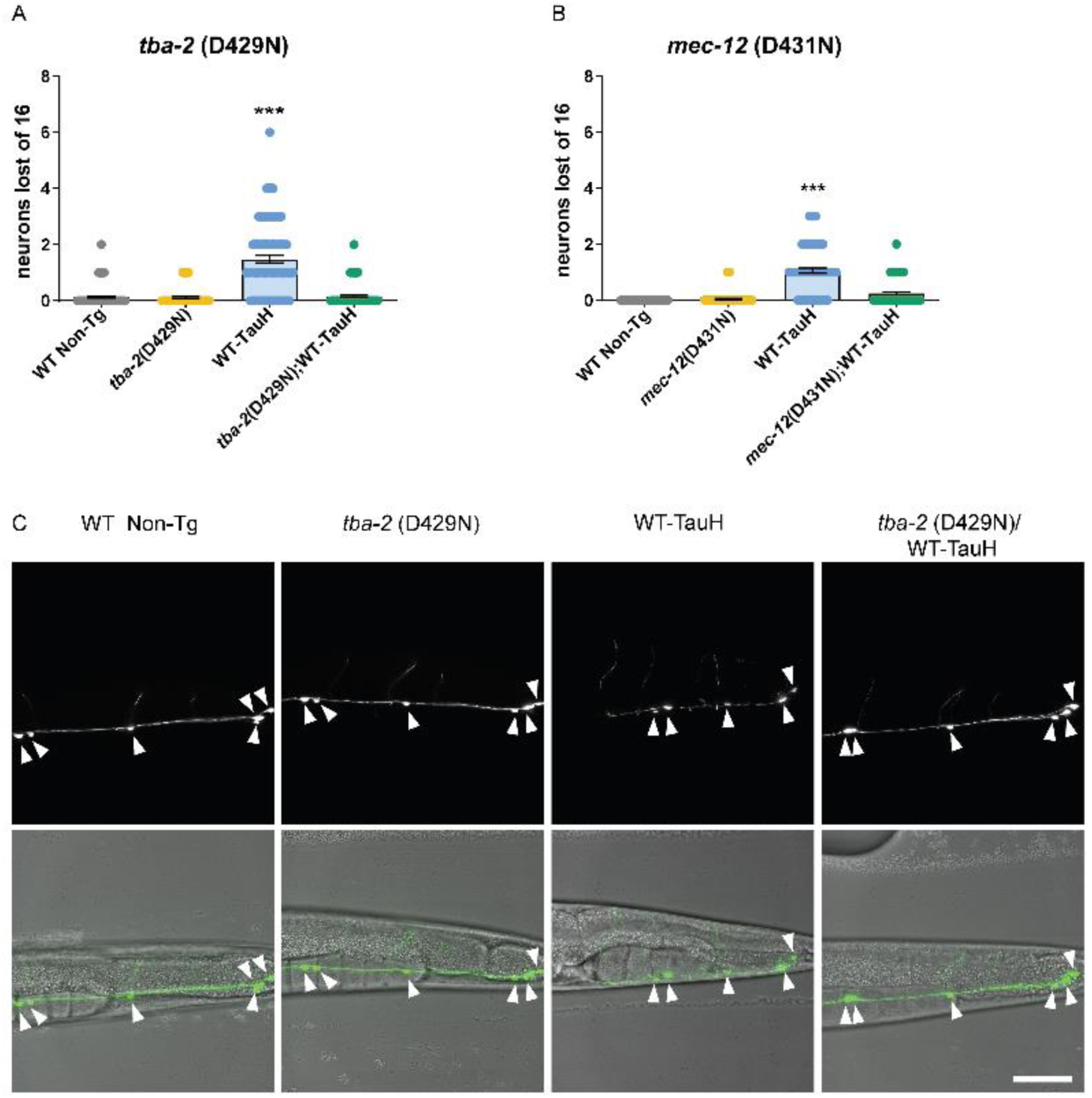
Mutant tubulins suppress neurodegeneration. *tba-2* (D429N) and *mec-12* (D431N) both suppress tau-induced neuron loss to wildtype levels (A) and (B) as visualized by GFP-labeled GABAergic ventral cord neurons (\*\*\**p<0.001,* n ≥ 82 animals error bars reflect SEM, Kruskal-Wallis test with Dunn’s multiple comparison post-hoc analysis). Representative images for *tba-2* (D429N)/WT-Tau and control animals (C). Scale bar (50µm).

### Mutant *tba-2* reduces levels of tau protein

To understand the molecular mechanisms underlying mutant-tubulin mediated suppression of tau toxicity, we assessed alterations to tau protein in mutant tau-transgenic strains versus tau-transgenic strains without tubulin mutations. We immunoblotted equivalent amounts of staged day-1 adult age matched tau-transgenic *C. elegans* using a phosphorylation and pan-isoform antibody to detect total tau levels across strains. We observed that *tba-2* moderately decreased overall tau levels by ∼45% while *mec-12* showed no significant effect on tau protein (**Figure 5 A-B**).

**Figure 5.**
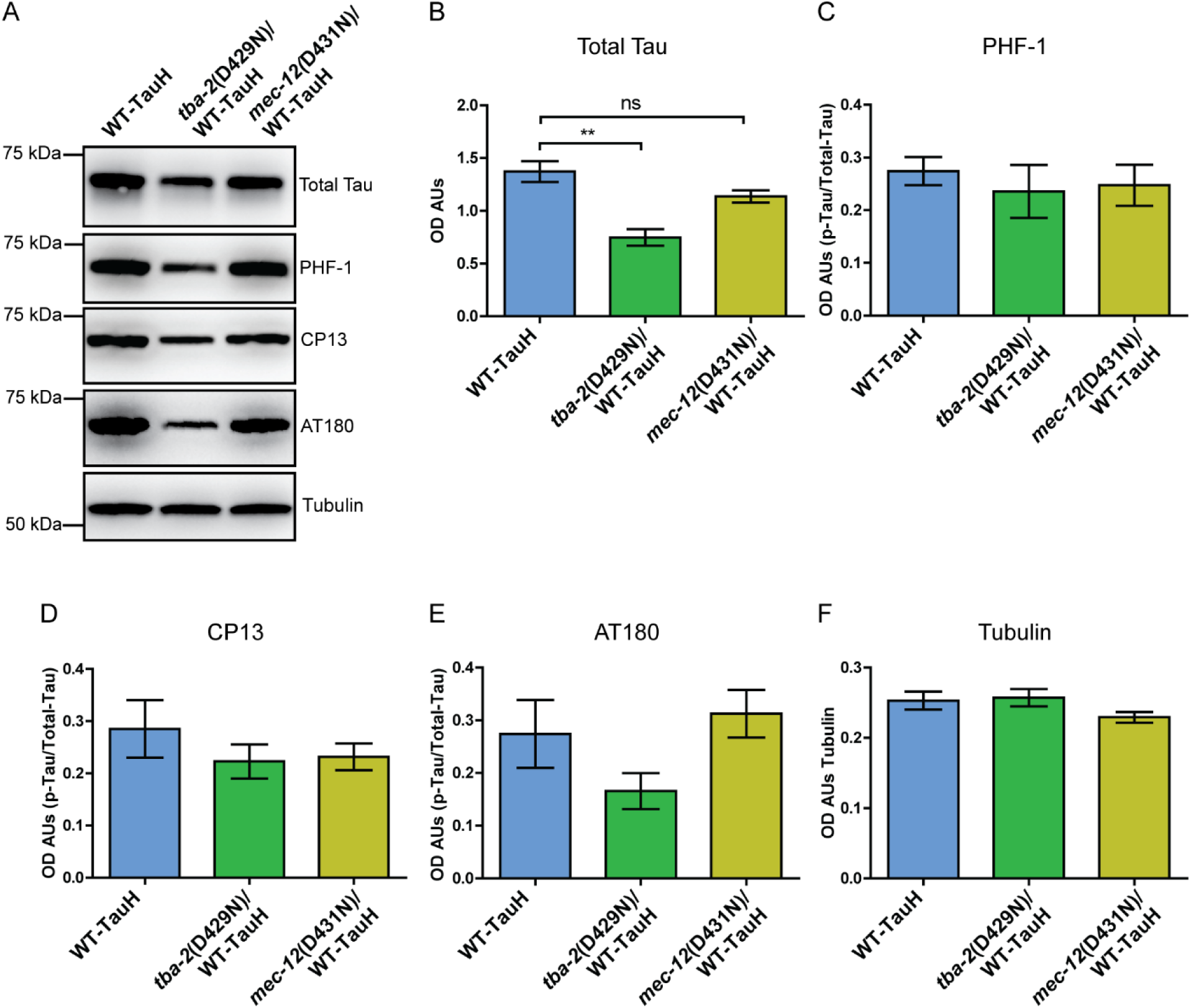
Tubulin mutations moderately alter total tau levels, without changing levels of tau phosphorylation. Western blot analysis of total protein from day 1 adult worms demonstrated that mutant *tba-2* (D429N) modestly but significantly reduced tau protein levels. However, significant reduction in tau was not observed in worms with mutant *mec-12* (D431N) (A and B). Tau phosphorylation was not altered significantly. (**p<0.01, 1-way ANOVA with Tukey post-hoc test, error bars reflect SEM). n = 4 populations (C-E, tubulin loading control in F).

Tau phosphorylation accumulates in neurons of AD affected brains as well as the neurons of *C. elegans* models of tauopathy. We, therefore, assessed for mutant-tubulin mediated alterations in tau phosphorylation. We immunoblotted staged day-1 adult *C. elegans* with antibodies specific to p-ser396/p-ser404 (PHF-1), p-ser202 (CP13), and p-thr231 (AT180) (**Figure 5A, C-E, antibodies described in Supplementary Table 3**). We observed that neither *tba-2* or *mec-12* altered levels of tau phosphorylation at these examined tau phospho-sites.

### Tubulin mutation does not influence tau aggregation

Progressive tau aggregation is a key pathological event in the progression of AD and related tauopathies and is recapitulated in *C. elegans* models. We hypothesized that mutant tubulin may confer suppression by preventing the aggregation of tau protein. We tested this hypothesis by extracting aggregated tau from whole *C. elegans* protein lysates using sequentially applied buffers of increasing solubilization strength to separate soluble from detergent insoluble tau species. The final solubilization buffer, formic acid (FA) fraction contains the most insoluble tau aggregates. Upon immunoblot analysis of the aggregated tau fractions, we detected no alteration of tau aggregation in tau transgenic *C. elegans* (WT-Tau, CK144) animals compared to *tba-2* mutants (**Figure 6**).

**Figure 6:**
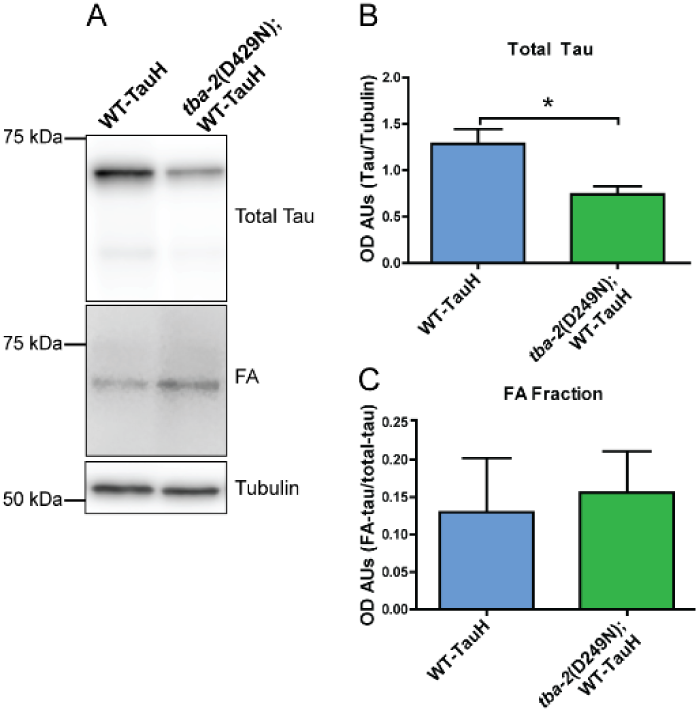
Mutant tubulin does not suppress tau aggregation in worms expressing high levels wildtype human tau. Western blot analysis of sequentially extracted fractions from day 1 so adult worm populations to isolate tau aggregates showed that mutant tubulin (*tba-2* (D429N)) reduced total tau levels but did not alter levels of tau aggregates (representative blots in A and quantitation in B, and C). FA shows formic acid soluble tau compared to total tau in whole worm lysates. (*\*p<0.05,* Student’s T-test, n = 4, error bars reflect SEM)

### Mutant *tba-2* decreases sedimented microtubule mass

Given that the tauopathy suppressing mutations occur in the region of the tubulin protein sequence that when folded and assembled into polymer, exists on the outer surface of the microtubule and within a predicted MAP interaction site, we hypothesized that the mutations may alter tubulin properties or function. To examine this possibility, we employed a microtubule sedimentation assay (based on [45], experimental schematic shown in **Supplementary Figure 6A**) in which *C. elegans* cellular extracts were used as the base material to build taxol stabilized microtubules. Taxol stabilized microtubules were isolated by ultracentrifugation allowing us to compare pelleted microtubule mass. (**Figure 7**). Signal from pelleted MT mass was normalized to corresponding input signal and mean post-incubation mass was calculated. We observed that *tba-*2/WT-TauH *C. elegans* extracts yielded less taxol-stable microtubule mass as compared to WT-TauH extracts *C. elegans* extracts (**Figure 7A, 7B**). Here we used nocodazole, a drug that prevents MT growth, as a negative, no microtubule, control (**Figure 7A**). Capillary western data from all 4 biological replicates is presented in **Supplementary Figure 6B.**

**Figure 7.**
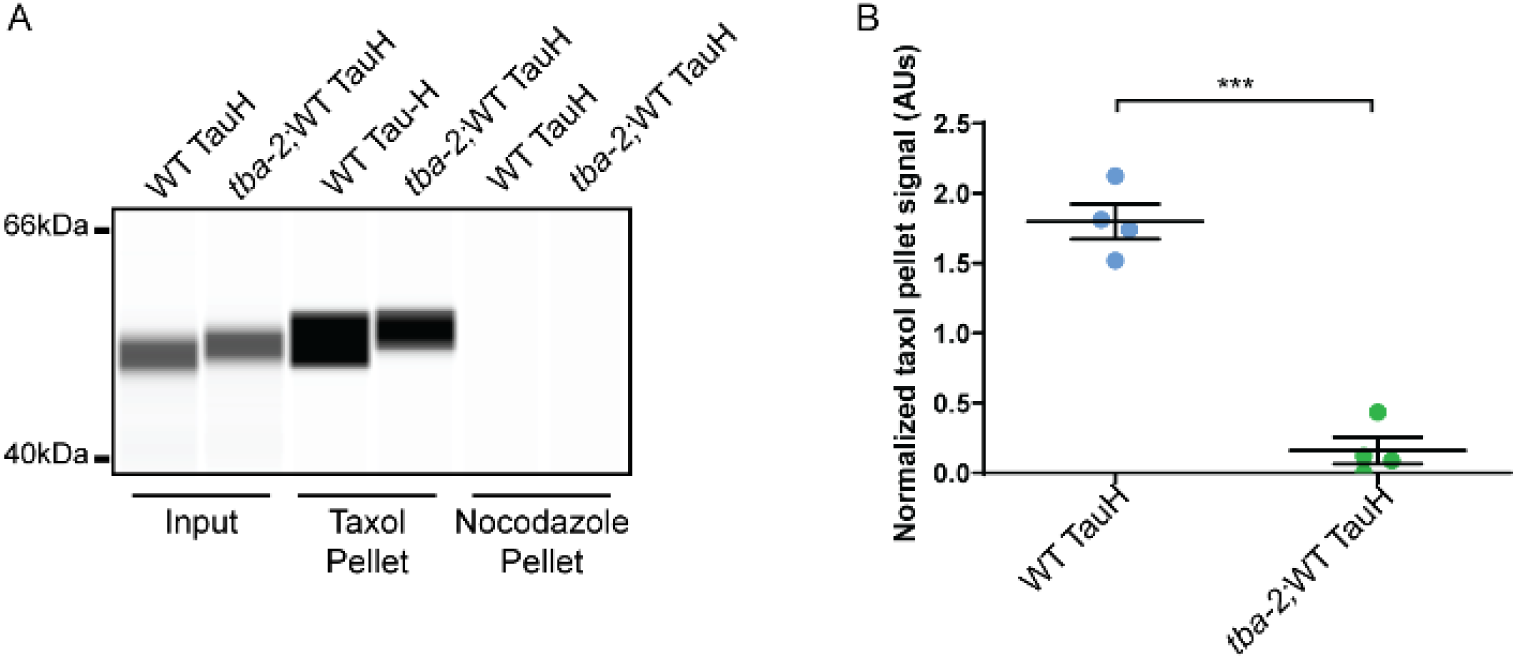
Mutation in *tba-2* changes taxol driven microtubule mass sedimentation. Capillary western performed using Peggy Sue^TM^ showing microtubule (MT) sedimentation resulted in reduced microtubule mass present after taxol incubation in the *tba-2*/WT TauH worm extracts as compared to extracts from WT TauH worms. No microtubule mass was detected with either extract when incubated with nocodazole (A). Data from 4 replicate capillary westerns were quantified in (B). Signal from taxol pellets was normalized to the corresponding pre-incubated input showing the reduction in sedimented MT mass seen with *tba-2*/WT TauH extracts was statistically significant *(***p < 0.0001*, unpaired T-test, n = 4, error bars represent SEM)

### Mutant tubulin does not alter the binding affinity between tau and soluble tubulin

The suppressor mutations map to helix 12 of α-tubulin on the outer lattice of the microtubule, which is predicted to be involved in tau-MT interaction. As such, we hypothesized that the mutations may confer suppression by altering the binding affinity between tau and tubulin. We tested this hypothesis using biolayer interferometry (BLI) to determine binding affinities (*K_d_*) between 6xHis-Tau(4R1N) (His-Tau) and wildtype and *tba-2* (D429N) tubulin association and dissociation measurements. Here, purified recombinant His-Tau was immobilized to NTA BLI sensors and introduced to solutions containing soluble tubulin isolated from wildtype and *tba-2* (D429N) *C. elegans* (tubulin association). After incubation in tubulin containing solutions, sensors were returned to buffer alone (BRB80), for tubulin dissociation. Changes to optical interference as detected through the sensor tips were used to calculate equilibrium association and dissociation constants. No significant difference in binding affinity was observed between the soluble tubulin isolated from wildtype animals as compared to *tba-2* (D429N) mutant *C. elegans* (**Figure 8**).

**Figure 8.**
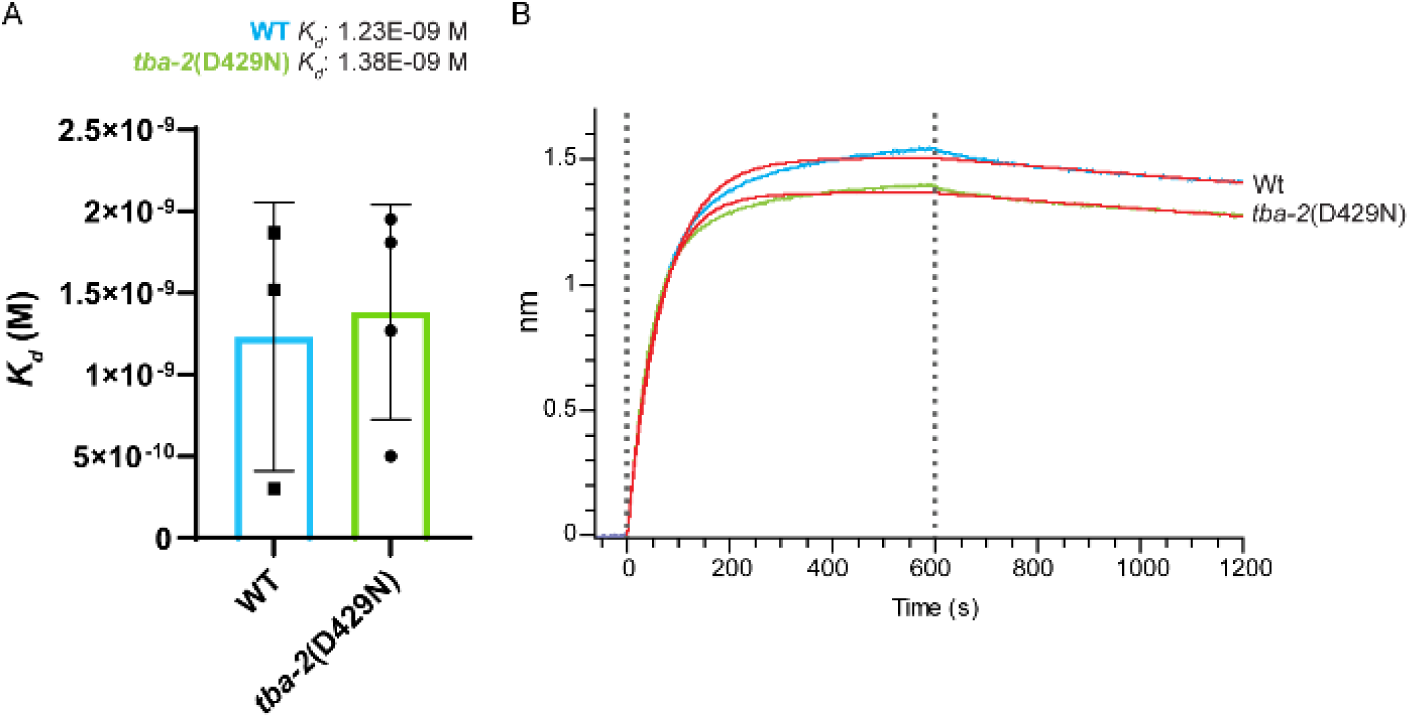
Mutant tubulin does not change binding affinity between Tau and soluble tubulin. Biolayer interferometry was used to determine the binding affinities between 6x-His tagged wildtype human 4R1N tau (His-Tau) and isolated wildtype (WT) or *tba-2*(D42N) *C. elegans* tubulin. The calculated *K_d_* between tau and *tba-2* (D429N) tubulin (1.38E-09 M) was not statistically different from tau-WT tubulin *K_d_* (1.23E-09 M) (A, Student’s T-test, n≥3, error bars represent standard deviation). Panel B shows representative binding curves for His-tau and WT tubulin (blue) versus His-tau and *tba-2*(D429N) (green), with corresponding theoretical fitted curves in red. **Supplementary Figure 7** shows representative acrylamide gels showing isolated *C. elegans* tubulin preparations.

## Discussion

Within the work presented here, we have shown mutations clustering in the tau-binding region of α-tubulin genes suppress tau-induced phenotypes in multiple *C. elegans* models of tauopathy, indicating the importance of this region in modulating microtubule function and protection from neurodegenerative assaults. The strongest suppressors identified completely rescue wildtype tau-induced phenotypes and strongly rescue mutant tau phenotypes and the phenotypes seen with tau/Aβ and Tau/TDP-43 co-pathology induced neuronal dysfunction. We have also identified similar mutations in other *C. elegans* α-tubulin genes that differentially suppress tauopathy to a level related to the relative level of tubulin expression from that gene; in other words, greater mutant tubulin expression confers a greater level of tauopathy suppression. Thus, titration of mutant tubulin levels may allow a fine-tuning of neuron specific tubulin expression to rescue tauopathy using a gene-based therapy approach.

All three mutant tubulin genes: the abundant mutant *tba-2,* the moderately abundant *tba-1* and the less abundant *mec-12* suppress the neurodegeneration seen in tau transgenic *C. elegans*. Both mutant *tba-2* and *mec-12* completely rescued neuron loss in tau transgenic animals to that of non-transgenic animals with minimal loss of GABA-ergic ventral nerve cord neurons. While these two mutations varied in their ability to suppress neuronal dysfunction, they are equally able to fully suppress neurodegeneration. Additionally, these data point towards neuron dysfunction as a major contributor to tau-induced motility phenotypes as compared to gross loss of neurons.

Mutant *tba-2* but not *mec-12* moderately reduces levels of total tau protein. The moderate reduction in total tau levels by *tba-2* and the absence of reduction with *mec-12* suggest that the mechanism underpinning the disproportionately remarkable rescue in neuronal dysfunction acts independently of tau levels. While we only see reduction in tau levels with the strongest suppressor, the strength of suppression seems outsized compared to the level of tau protein reduction and points to a mechanism independent of total tau levels. Additionally, we observed no alteration to phosphorylation or aggregation of tau with mutant *tba-2,* suggesting these mutants to confer suppression through a mechanism independent of pathological tau accumulation.

The amino acid substitutions that confer tauopathy suppression occur in or immediately adjacent to helix 12 of α-tubulin. Helix 12 is positioned on the outer surface of the microtubule once tubulin subunits are assembled into the polymer. Since helix 12 is important for MAP interactions we hypothesized that the mutations might affect microtubule properties in assembled microtubules. We tested this by forming MTs from endogenous *C. elegans* tubulin with the stabilizing drug, taxol, and measured the relative microtubule mass pelleted upon ultracentrifugation. Interestingly, we found that less microtubule mass pelleted from lysates that contained mutant tubulin, indicating that the microtubules containing mutant tubulin are either taxol resistant or have different assembly properties in the presence of taxol. Microtubule stabilizing compounds have previously demonstrated rescue of tau pathology in mammalian models improving axon density, cognitive function and clearance of tau pathology [26, 27]. Taken together, these data support the notion that microtubules containing mutant α-tubulin possess different properties as compared to those made solely of wild type tubulin and altering microtubule properties may contribute to increased functionality in the presence of pathological tau.

The cluster of tau suppressing mutations occur in helix 12, a region playing a prominent role in the charge-charge interactions facilitating tau-MT binding. To test the hypothesis that mutations in this region may alter tau-MT binding we used recombinant human tau immobilized to the biolayer interferometry (BLI) sensor. Using solutions containing tubulin isolated from wildtype tubulin or *tba-2*(D429) *C. elegans* we could determine association and dissociation parameters for each. The binding affinities between tau and soluble wild type tubulin compared to tau and mutant tubulin were not observably different. However, this does not rule out a difference in tau binding to polymerized microtubules containing mutant tubulin. Additional testing will be needed in order to explore whether mutations affect tau or other MAP binding with polymerized MTs.

### Conclusions

While more experimentation is needed to understand the specific molecular details of tauopathy suppression, it is already evident that the mutations described in this study provide protection from the adverse effects of tau-mediated neuronal dysfunction and neurodegeneration. As the mutations do not reduce the level of phospho-tau or tau aggregates, it seems the α-tubulin mutations do not act by directly reducing pathological tau species but rather protect neurons via a tubulin/microtubule function mediated mechanism. Given recent advances in gene therapy technologies for neurodegenerative and neurodevelopmental disorders [46, 47], the possibility of harnessing mutant helix 12 α-tubulin to suppress tauopathy phenotypes appears attractive. Therefore, exploration of the mechanism of action of these mutations will contribute to a more complete understanding of the conceptual basis for developing effective disease modifying therapeutics for tauopathy.

## Materials and Methods

### *C elegans* strains and maintenance

For a list of strains used, refer to **Supplementary Table 2**. Worms were maintained at 20°C (with the exception of Aβ_1-42_ strains and controls, which were maintained at 15°C prior to temperature upshift for Aβ_1-42_ peptide expression for 24 hours at 25°C prior to experimentation, on NGM plates seeded with OP50 *E. coli* as described in [48]. The *C. elegans* tubulin alleles described in Supplementary Table 1 were generated through chemical mutagenesis using ethyl nitroso urea as the mutagen using established methods previously described in detail [34, 35, 49].

For mRNA and protein extraction, worms were grown on nutrient rich media plates containing 5 times peptone (5xPEP) seeded with OP50 [48]. For microtubule co-sedimentation experiments, worms were grown to dense populations on 5xPEP plates and subsequently washed onto egg plates which were prepared by seeding OP50 mixed with separated chicken egg yolk. Animals were subsequently harvested from 5xPEP plates using sequential washes with M9, or by floating on sucrose cushion if harvested from egg plates [32, 50, 51]. Pellets were snap frozen in liquid nitrogen and stored at −80°C until use.

### *C. elegans* motility assays

*C. elegans* were synchronized and grown to day 1 of adulthood on NGM growth plates. Plates containing the worms were placed in the assay room and allowed to acclimate for 20 minutes, prior to assay initiation. One strain at a time, worms were washed from NGM plates to food free video plates with 2mL of M9 and allowed to acclimate to the buffer for 10 seconds before a 1 minute video was recorded. Video data was collected using the WormLab platform (MBF BioScience, VT). Worm locomotor behavior was tracked and analyzed using the WormTracker software (MBF BioScience, versions 2019, 2020 and 2022). Body bends from the mid-point body location of each worm were counted. The total number of body bends was divided by the track length(s) to give the frequency of body bends per second, this was then converted to body bends per minute. Mean body bends per minute was compared across strains using the Kruskal-Wallis test with post-hoc Dunn’s test.

### Manual thrashing assay

A manual thrashing assay [50] was performed to assess the efficacy of suppression of a single copy of the *tba-2*(D429N) mutation. *tba-2* (D429N)/+ heterozygous strains were generated by mating the homozygous *tba-2;* WT-Tau strain with WT-Tau males to make obligate heterozygotes for *tba-2(D429N)* in the F1 generation. F1s were subjected to a manual thrashing assay where worms were placed in a 10 µL drop of M9 buffer on a microscope slide. After worms were allowed to acclimate for 10 seconds, the number of thrashes were counted for 1 minute. One thrash is defined as a bend to the opposite direction across the midline and a return to starting position or two complete bends to the same side where the worm straightens to the midline in between. To confirm heterozygosity worms were removed from the buffer and placed on a fresh NGM plate and a clonal population was allowed to grow from the single worm. The populations were then genotyped for *tba-2(D429N)* heterozygosity.

### Quantification of mRNA levels of tubulin *sut* genes in *C. elegans*

*C. elegans* were grown from eggs to day 1 of adulthood at 20°C on 5xPEP plates. Worms were washed off plates using M9 buffer and pelleted by centrifugation to a final pellet size of ∼100mg. Worm pellets were snap frozen in liquid nitrogen and stored at −80°C. RNA was isolated from ∼100mg pellets using TRIzol Reagent (Life Technologies) according to the manufacturer’s instructions. RNA was quantified using a NanoPhotometer NP80 (Implen GmbH, Munich, Germany) and integrity was assessed by 1% Tris/Borate/EDTA (TBE) agarose gel electrophoresis. RNA samples were treated with DNA-*free* DNAse treatment and removal kit (Invitrogen) following the manufacturer’s instructions. cDNA was made using the iScript Reverse Transcription Supermix (Bio-Rad) following manufacturer’s instructions and qPCR was performed iTaq Universal SYBR Green Supermix (Bio-Rad) in a CFX Connect Real-Time PCR System (Bio-Rad). Data was analyzed using CFX Manager 3.0 (Bio-Rad). Experimental samples were run in duplicate and three independent biological replicates were performed. Total *tba-2* and transgenic *tba-*2 were normalized to *rpl-32* internal control gene. Means were compared using a Student’s T Test. Primers used were as follows: Transgenic-*tba-2* forward: GAC TCC AAC GAG GGA GGA G, Transgenic-*tba-2* Reverse: CGA GAT GGC GAT CTG ATG, Total-*tba-2* Forward: TCT CGT CTC GAC TAC AAG TTC G, Total-*tba-2* Reverse: CTT CTC GAG AGC AGC CAA GT, *rpl-32* Forward: GGTCGTCAAGAAGAAGCTCACCAA [50], *rpl-32* Reverse: TCTGCGGACACGGTTATCAATTCC [50].

### GABAergic Neuron Loss Assay

GABAergic neuron loss was assessed as a measure of gross neurodegeneration [32, 44, 50]. Tubulin mutant;Tau-transgenic worms were crossed into CZ1200 (juIs76 [unc-25p::GFP + lin-15(+)]) to generate *tba-2*;WT-Tau;unc-25p::GFP, WT-Tau;unc-25p::GFP, *tba-2*;unc-25p::GFP, *mec-12*;WT-Tau;unc-25p::GFP and *mec-12*;WT-tau;unc-25p::GFP. Worms were synchronized to day 1 of adulthood (approximately 4 days post-hatch) and mounted onto 2% agarose pads and immobilized with sodium azide. The number of visible GABAergic neurons along the ventral nerve cord were counted using a DeltaVision microscope (Applied Precision) at 60x magnification. A minimum of 10 animals were counted per strain per replicate, and a minimum of 3 biological replicates were completed. Statistics were calculated using the Kruskal-Wallis test with a post-hoc Dunn’s comparison, \*\*\**p<0.001,* error bars reflect SEM.

### Protein extraction for total tau levels

Protein extraction and immunoblotting procedures were conducted as previously described in [29, 32]. Worms were grown from eggs to staged day 1 adult populations on 5xPEP plates. Worms were harvested by washing with M9 buffer, and pelleted by centrifugation (3,000xg, 45 sec). Pellets were resuspended and washed three times with 8mL of M9 buffer and aliquoted to approximately ∼100mg final pellet size into pre-weighed Eppendorf tubes. Buffer was aspirated from centrifuged worms and pellets snap frozen in liquid nitrogen prior to storage at −80°C. Whole worm protein lysates were generated as follows. Worm pellets were weighed to determine pellet mass and thawed on ice. SDS proteins ample buffer (0.046 M Tris, 0.005 M ethylenediamine tetraacetate, 0.2 M dithiothreitol, 50% sucrose, 5% sodium dodecyl sulfate, 0.05% bromophenol blue, 1× concentration) was added to the pellets at a volume (µL) four times the pellet weight (mg). Pellets were sonicated three times for 20 seconds each at 70% amplitude, and cooled ice between sonication sessions. Samples were boiled for 10 min at 95°C and centrifuged at 13,000xg for 2 minutes prior to gel loading. Four biological replicates were performed.

### Sequential protein extraction

Insoluble tau aggregates were isolated from soluble tau by the collection of sequential fractions after worm lysate treatment with buffers of increasing protein solubilization strength, as previously described in [29]. Briefly, staged day 1 adult worm pellets were resuspended in 2µL per milligram of wet pellet weight of RAB high salt buffer (0.1 M MES, 1 mm EGTA, 0.5 mm MgSO4, 0.75 M NaCl, 0.02 M NaF, pH 7.0, plus Complete Protease Inhibitor Cocktail (Roche Diagnostics) and 0.5 mm PMSF). Samples were sonicated three times at 70% amplitude for 8 seconds. An appropriate amount of worm lysate was reserved as the total fraction used to determine the relative level of total tau (both soluble and insoluble), while the remaining RAB sample (minimum volume of 100 µL) was subjected to centrifugation at 40,000xg for 40 min. The supernatants (RAB soluble tau) were collected and pellets resuspended in RIPA buffer (150 mm NaCl/1% Nonidet P-40/0.5% deoxycholate/0.1% SDS/50 mm Tris, pH 8.0, plus Complete Protease Inhibitor Cocktail (Roche Diagnostics) and 0.5 mm PMSF), using a volume of RIPA buffer equal to half that of the RAB sample starting volume. Samples were sonicated 1-2 times at 70% amplitude for 8 seconds and subsequently centrifuged at 40,000xg for 20 min. The supernatants were collected as the RIPA-soluble fractions (detergent-extractable insoluble tau). The remaining pellets were resuspended in a volume of 70% formic acid (FA, in water), equal to the volume of the starting RAB sample. FA resuspended pellets were sonicated 1-2 times at 70% amplitude for 8 s and centrifuged under vacuum in a SpeedVac with no additional heat until FA had evaporated. The remaining pellets (FA insoluble tau) were resuspended in a volume of 2x SDS protein sample buffer (0.046 M Tris, 0.005 M ethylenediamine tetraacetate, 0.2 M dithiothreitol, 50% sucrose, 5% sodium dodecyl sulfate, 0.05% bromophenol blue, 2x concentration). FA samples were then sonicated 1-2 times at 70% amplitude for 8 s. Samples were then boiled for 5 minutes (95°C) and centrifuged at 13,000xg at 4°C for 5 minutes. RAB samples were boiled for 5 min (95°C), chilled on ice for 5 min then centrifuged at 13,000xg at 4°C for 15 minutes and supernatants were collected as RAB-soluble fractions. SDS protein sample buffer was added to a final concentration of 1x to total, RAB-soluble and RIPA-soluble fractions, samples were boiled for 5 min at 95°C, chilled on ice for 5 min, then centrifuged at 13,000xg for 5 min. All samples were stored at −20°C prior to SDS-PAGE electrophoresis. Four biological replicates were performed.

### Immunoblotting

Lysed total worm protein or sequentially extracted samples were subjected to SDS-PAGE using precast Tris-HCl gels (catalog number 3450028, BioRad, Hercules, CA), according to manufacturer’s instructions. Gels were run at 200V for approximately 60 minutes and proteins were cold transferred to PVDF membranes at 80V for 30 minutes. Membranes were blocked with 5% milk in PBST (1x PBS with 0.1% Tween-20) for 2 hours and incubated in primary antibody solutions diluted in 5% milk PBST block overnight at 4°C. Membranes were washed with PBST three times for 10 minutes each wash. Membranes were then incubated in secondary antibody solution for 2 hours at room temperature (see Supplementary Table 3 for a list of antibodies and dilutions used), after which they were washed three times in PBST as above. Detection was completed using the Clarity Western ECL substrate kit (Biorad; cat. number 1705060) and LiCOR Odyssey and Image Studio software. Blots were quantitated using ImageJ [52].

### Large Scale growth of *C. elegans* using chicken egg yolk growth media

Worms were grown at large scale on 5X Peptone (PEP) plates seeded with an egg yolk-OP50 mixture based on methods previously described [51, 53]. Briefly, approximately 1 egg per 5X PEP plate to be seeded was sterilized in a 95% ethanol bath for approximately 5 minutes. The eggs were removed and air dried on a sterilized bench for several minutes. Egg yolks were separated using standard culinary methods maintaining as much sterility as possible (cracked on the side of an autoclaved beaker, and a sterile egg culinary egg separator was used). Separated egg yolks were placed in an autoclaved beaker. Using a sterile kitchen fork, as much non-yolk material and debris was removed as possible. Alternatively, equivalent volume of store-bought pasteurized separated egg yolks were used. The remaining yolk was mixed with 1.5-2mL stationary phase OP50 bacteria grown overnight at 37°C in standard LB. The egg-yolk/OP50 mixture was sterile pipetted onto 25cm 5X PEP plates, using 12mL per plate. Each plate was swirled to spread the egg-yolk/OP50 mixture and additional debris was pipetted off. Plates were left for approximately 3 days at room temperature to solidify to custard like consistency at which point they were seeded with *C. elegans*. *C. elegans* were grown to a dense lawn and harvested by sucrose floatation.

### Sucrose Flotation to clean *C. elegans* cultures

As described previously in [51], worms from egg plates were harvested using room temperature M9 and collected in 50mL conical vials. Worms were pelleted by centrifugation 2500xg at 4°C in an Eppendorf tabletop centrifuge and resuspended in cold fresh M9 to rinse, and re-pelleted. Supernatant was discarded and pellets were resuspended and brought to 25mL with cold M9 and incubated on ice for ∼10 minutes. 25mL of cold 60% sucrose was added. Tubes containing the worm and sucrose mixtures were inverted 10 times to mix. They were then centrifuged for 5 min at 4°C at 2500xg. Worms float and form a supernatant layer on sucrose whereas bacteria and debris pellet at the bottom of the tube. Worms were removed with a sterile metal spatula and wide bore 1mL micropipette into fresh cold M9 on ice. Total volume was brought up to 50mL with cold M9. Tubes were inverted to mix thoroughly and centrifuged for 2 min at 1000xg at 4°C. The supernatant was discarded and fresh cold M9 was added to 30mL volume to resuspend the pellet and rinse any residual sucrose. Worms were centrifuged a final time at 1000xg for 2 min at 4°C. The supernatant was discarded and the pellet snap frozen in liquid nitrogen then stored at −80°C until use.

### Microtubule Sedimentation assay

MT sedimentation assays were based on the protocol published in [45]. Briefly, large scale populations were grown on plates with chicken egg yolk media and sucrose floated to clean. Approximately one gram worm pellets were weighed and thawed on ice. Worm Lysis Buffer (50mM HEPES-KOH, pH 7.6, 1mM MgCl2, 1mM EGTA, 75mM KCl, 0.5mM DTT plus ProteincOmplete™, Mini, EDTA-free Protease Inhibitor Cocktail) volume (mL) was added to 1.5 times the weight (g) of the pellet. The pellets were resuspended and sonicated at 70% amplitude for two rounds of 30 seconds on 60 seconds off, for a 1 minute sonication time per round that was repeated for two rounds to complete 2 minutes of active sonication time. Samples were kept on ice throughout the sonication process and the sonicator tip was cooled with ice bath in between each sonication duration. Crude lysates were ultra centrifuged 20,000xg for 10 minutes (4°C). The Low Speed Spin (LSS) supernatant was saved and subjected to centrifugation at 50,000xg (4°C). The supernatant from this spin was reserved as the High Speed Spin (HSS) sample, this material was snap frozen with liquid nitrogen and stored at −80 until use. HSS material was thawed on ice prior to conducting microtubule sedimentation assays. The total protein concentration for the extracts was determined by Bradford assay and equalized prior to proceeding with the sedimentation reactions. DTT and GTP was spiked into 1mL of the equalized HSS sample (final concentrations of 0.5mM and 1mM respectively) to make the HSS++ input material used in the subsequent reactions. Material was kept on ice until reactions were assembled. Three reaction conditions were assembled using 205µL HSS++ in each reaction: DMSO control, Nocodazole control and Taxol stabilized. DMSO was added to the equivalent volume as the additions of nocodazole (10µg/mL final concentration) and taxol (20µM final concentration). Microtubule reactions were incubated at 25°C for 10 min to allow for microtubule assembly. 200µL of the reactions was placed on a warmed (25C) glycerol cushion (40% glycerol, 50mM HEPES-KOH, 1mM MgCl2, 1mM EGTA, 75mM KCl, 0.5 mM DTT and ProteincOmplete™, Mini, EDTA-free Protease Inhibitor Cocktail). Microtubules were spun over the glycerol cushions at 48,000xg for 30 minutes (25°C). 80 µL of the supernatant was collected and combined with 20 µL 5x SDS sample buffer. The supernatant-cushion interface was washed 3 times with 200 µL worm lysis buffer. The cushion was carefully removed by pipetting and the residual pellet was resuspended in 200 µL 1x SDS sample buffer. Samples were then subjected to capillary electrophoretic analysis using the Simple Western Peggy Sue^TM^ Capillary Electrophoresis System for analysis of tau and tubulin.

### Capillary electrophoresis and immunodetection

We used the Peggy Sue^TM^ (ProteinSimple) Simple Western system to immunoblot MT sedimentation assay samples following the manufacturer’s instructions. Briefly, samples were diluted with 0.1X sample buffer and 5× Fluorescent Master Mix, boiled for 5 minutes at 95°C, cooled and loaded into a 384-well assay plate (12-230kDa Size Separation module, Catalog No. SM-S001, ProteinSimple). Samples were immunoblotted using the E7 (1:100) β-tubulin primary antibody, SP70 (1:2000) and DAKO (1:10,000) total tau primary antibodies, and goat anti-mouse-HRP (1:100) and goat anti-rabbit (1:100) secondary antibodies (**Supplementary Table 3**) that were diluted in Antibody Diluent 2. Samples were run and analyzed using the default assay in Compass for SW Version 4.0.0 (ProteinSimple). Signal for sedimented MT mass was normalized to total pellet signal then compared to input signal (e.g. taxol pellet WT/(taxol-pellet(WT) + taxol-(*tba-2*))/input tubulin signal) and means were calculated using four biological replicates. A Student’s T Test was used to determine the statistical significance of the difference between the means.

### Isolation of *C. elegans* tubulin

Tubulin isolation methods were based on those previously reported in Widlund et al. 2012, [54]. Large scale populations were grown on plates containing chicken egg yolk media and sucrose floated to clean. Approximately 10g sucrose floated worm pellet material was thawed, mixed with equivalent volume (weight to volume) of BRB80 buffer (80mM PIPES, 1mM EGTA, 1mM MgSO_4_, pH 6.8) to form a slurry. Slurry was re-frozen dropwise in liquid nitrogen to make pellets. Pellets were enclosed in a folded foil envelope, ∼15ml pellets per batch, and hammered over a brick of dry ice. Once pellets were ground into fine powder, powder was transferred to pre-chilled 50ml conical on dry ice. BRB80 was added to bring the volume to 50mL, adding more BRB80 as the worm powder thawed to maintain the 50mL volume. The crude lysate was subjected to a low speed spin at 3500 x g at 4°C. The supernatant was decanted into a fresh 50mL conical and pellet discarded. The supernatant was subjected to high-speed ultracentrifugation at 301,311 x g (50,000 rpm in Beckman Ti50.2 rotor) for 35 minutes at 4°C. The centrifuged supernatant was then filtered through a cell strainer (Falcon 70µM, catalog number 352350) 0.22µM filter to further clear lysate. Cleared lysate was loaded, by gravity flow, onto an affinity column with TOG1/2 domains from the *Saccharomyces cerevisiae* protein Stu2. Column was washed with 10 column volumes of wash buffer comprised of (1x BRB80, 100uM GTP and 100uM MgCl2). Tubulin was eluted off the column using 3 column volumes of elution buffer (1xBRB80, 10uM Mg2 GTP, and 500mM (NH4)2SO4). Protein containing fractions were determined using Bradford dye (10µl fraction sample to 100µl diluted Bradford reagent, Biorad catalog number 5000006) to assess protein presence by eye. Protein containing fractions were pooled, mixed and desalted over BRB80 equilibrated PD10 desalting columns (Sigma, catalog number GE17-0851-01). Desalted protein was concentrated to 250µl at 4°C at 3500 x g with 30 kDa molecular weight cut off Amicon Ultra centrifugation filters (Millipore catalog number UFC9030) previously equilibrated with BRB80. Glycerol was added to a final concentration of 5% v/v and mixed thoroughly. Tubulin was aliquoted and snap frozen in liquid nitrogen and stored at −80°C until use.

### TOG1/2-column for affinity purification of *C. elegans* tubulin

Purification of GST tagged Stu2 TOG1/2 domains and affinity column creation was based on methods in Widlund et al [54]. A 90 L growth and induction of pGEX-6P-1 Stu2 1-590 (Addgene plasmid #38314) was expressed in Rosetta (DE3) cells and bacterial cell paste was frozen into a brick and stored at −80°C until use. At the time of TOG1/2 protein isolation, chunks of the frozen *E coli* cell paste (∼20g) were thawed on ice. Bacteria was resuspended in 140mL of lysis buffer (2xPBS, 12.5mM DTT, and cOmplete Mini-EDTA free protease inhibitor pellets (Millipore, catalog number 11836170001). 0.1g lysozyme, 100uL DNAse, 25uL RNAse A, 80uL 1M MgCl2 was added to bacterial resuspension to initiate lysis. Resuspension was inverted to mixed every ∼5 min for 20 minutes incubating on ice. Resuspension was aliquoted into 4 conicals and sonicated for one round at 70% amplitude for 10 seconds. Lysate was then transferred to oakridge tubes for centrifugation in a Sorvall SA-600 rotor at 16,000 RPM at 4°C for 30 minutes. Cleared lysate was filtered through a 70µm cell strainer (Falcon, Fisher Scientific catalog number 352350) then passed through a 0.22µM filter to further clear. Lysate was applied to a 5mL GSTrap 4B Sepharose column (Cytiva 28401748) connected to a peristaltic pump and equilibrated with equilibration buffer (2xPBS, 1mM DTT) at a flow rate of 2mL per minute. The column was washed with 5 column volumes (CV) of wash buffer 1 (W1, 2x PBS, 0.1% Tween20, 1mM DTT) followed by 2.5 CV of wash buffer 2 (W2, 6X PBS, 1mM DTT), 2.5 CV of wash buffer 3 (W3, 2x PBS, 1mM DTT). Protein was eluted with Elution buffer (2x PBS, 50mM glutathione, 10mM DTT, pH 9.0) collecting 1 ml fractions. Protein presence was detected using a small-scale Bradford assay as described above. Protein containing fractions were desalted using PD10 desalting columns equilibrated with pH 10 coupling buffer (0.1M sodium citrate, 0.05M Sodium carbonate, pH brought to 10 with NaOH). Protein was snap frozen in liquid nitrogen and stored at −80°C until used for coupling to resin.

Resin coupling was performed using the AminoLink™ Plus Immobilization Kit, 2 mL format (Thermo Scientific, 44894). Approximately 40mL volume and 50g of Stu2 TOG1/2 protein was thawed and mixed together, resin from two columns (4mL) was washed using pH10 coupling buffer and using coupling buffer, resuspended and transferred to with the thawed Stu2 TOG1/2. Resin was incubated in the protein solution with end over end rotation for 1 hour at room temperature. Coupled resin was divided and packed into two columns. Resin was then washed twice with 2mL of pH 10 coupling buffer. Each column was then washed with 10mL of 1xPBS followed by 3 washes with 2mL of storage buffer (1xPBS, 50% v/v glycerol). After the final wash in storage buffer, additional storage buffer was used to top off the column head space, the top and bottom caps were securely replaced, columns parafilmed to seal and placed upright at −20°C for storage. Columns were dedicated either to wildtype or *tba-2*(D429N) tubulin isolations and re-used for at least 5 isolations.

### Isolation of 6X-histidine tagged recombinant human 4R1N tau protein

E. coli codon-optimized 6X-histidine-tagged human 4R1N tau (His-tau) cDNA was cloned into the pET-28a vector (Merck KGaA, Darmstadt, Germany) and transformed into BL21 (DE3) bacteria (New England Biolabs, Ipswich, MA). Ten mL starter cultures of Terrific Broth (TB) were grown overnight at 37°C in a shaking incubator. For induction, 1 L cultures of TB were inoculated, grown with shaking at 37°C to log phase, and induced with 1mM IPTG for 3 hours at 37°C. Bacterial pellets were resuspended in lysis buffer (1X PBS supplemented with 150mM NaCl, 5mM β-mercaptoethanol) and boiled to lyse. Lysate was cooled in ice-bath before centrifuging at 7000 x g to clear lysate. Cleared lysate was incubated with Ni-NTA agarose resin (Qiagen, Hilden, Germany) which was washed once with 10X bed volume of lysis buffer and then twice using lysis buffer with 10mM imidazole. His-tau was eluted twice: first, using 1 bed volume of lysis buffer with 250 mM imidazole and then using lysis buffer with 500 mM imidazole. Elution 1 was desalted once with PD10 columns (Cytiva, Marlborough, MA) into 50 mM HEPES, 25 mM NaCl, pH 7.4 and Elution 2 was desalted twice.

### Biolayer interferometry assay

Wildtype and *tba-2*(D429N) tubulin was isolated as described above. Recombinant wildtype human 4R1N 6xHis-tagged tau protein (His-tau) was prepared as previously published in McMillan et al 2023 [55]. Recombinant human His-tagged MPRO protein (His-MPRO) was prepared as previously published in Baker et al 2021 [56].y His-MPRO was used as both a quencher and a control for non-specific binding.

All data was obtained using a Sartorius Octet R8 with Sartorius Octet NTA Biosensors (18-5101 Lot number 2412000511) in a 96-well format. BRB80 buffer (80mM Pipes, 1mM EGTA, 1mM MgSO4, pH 6.8) was used to establish an initial baseline. After a 10 minutes shaking (1000 RPM), experimental steps proceeded at 30°C shaking at 1000 RPM in the following order: loading of His-tau (75 nM), quenching with His-MPRO (10 ug/ml), washing with BRB80, running a secondary baseline with BRB80, association, and lastly dissociation with BRB80. After a dissociation step with BRB80, the sensors were regenerated with 10 mM glycine (pH 1.5 – 2.0), reactivated with 10 mM NiCl, and washed with BRB80. Washes, quenches, activation, and loading steps ran for 300 seconds each. Association and dissociation steps ran for 600 seconds each. Baseline steps ran for 180 seconds each. The regeneration step ran for 30 seconds. Shaking was maintained at a speed of 1000 RPM throughout the experiment. Three to four independent isolations, for each wildtype and *tba-2* (D429N) tubulin were tested at multiple concentrations within 133 −384.4 nM range. His-MPRO (75nM) was used in place of His-Tau as a test for non-specific binding by tubulin and BRB80 was used as a buffer only control.

Analysis was conducted in the Octet Analysis Studio Software (v. 13.0.0.32). Background signal was accounted for with the His-MPRO control sensor and the buffer only initial sample within the software. Graphs were aligned on the Y-axis according to the baseline averages, along with an inter-step correction aligned to the dissociation step and filtered via Savitzky-Golay. Data was then exported into Microsoft Excel. The mean of the dissociation coefficients (*K_d_*) for a given tubulin preparation was calculated to determine the *K_d_* for that tubulin isolation. The means of the wildtype or *tba-2* (D429N) isolations were calculated to compare the overall wildtype-tau *K_d_* versus mutant-tau *K_d_*. Isolation replicate means were graphed in Graphpad Prism 10. Student’s T test was used to test the difference in mean *K_d_* and the displayed association curves are representative curves for wildtype and mutant worm tubulin obtained in the same replicate.

## List of abbreviations

FTDP-17: Frontotemporal Lobar Dementia with Parkinsonism chromosome 17 type
AD: Alzheimer’s Disease
MT(s): microtubule(s)
MAPs: Microtubule Associated Proteins
TDP-43: TAR DNA-binding protein 43
sut: suppressor of tau Pathology
CeNGEN: *C. elegans* Gene Expression Network
Aβ: Amyloid-β
His: Histidine
WT: wildtype
Tg: transgenic

## Declarations

### Ethics Approval and Consent to Participate

NA

### Consent for publication

NA

### Availability of data and research materials

Data generated in this work is available at Dryad data repository https://doi.org/10.5061/dryad.15dv41p9s

Reagents and materials generated in this study will be made available from the corresponding author upon reasonable request to corresponding author.

### Author contributions

SB conceptualized, designed and executed experiments, analyzed data, drafted manuscript. AS, MB, RU, JS, KK and CD created reagents (i.e. *C. elegans* strains, purified protein) and executed experiments, RK performed forward genetic screening. LW, NL and BK aided in conceptualization, experiment design and design of analyses. All authors read and edited manuscript.

## Acknowledgments

We thank Charles L. Asbury for guidance and mentorship in producing this work and thank Asia Beale for efforts in growing large populations of *C. elegans* for tubulin isolations and sedimentation assays. The work presented here is the responsibility of the authors and does not represent the official views of the NIH or government of the United States of America.

## SUPPLEMENTARY MATERIAL

**Supplementary Table 1.** Mutant tubulin Alleles identified in forward genetic screening.

| Gene name | Strain name | AA change | Codon change |
| --- | --- | --- | --- |
| *mec-12* | CK731 | D431N | GAT-->AAT |
| *tba-1* | Ck1753, CK2050 | E437K | GAG-->AAG |
|  | CK2051 | A429V | GCT-->GTT |
|  | CK2192 | E436K | GAA-->AAA |
|  | CK4295 | R425H | CGT-->CAT |
| *tba-2* | CK1458, CK1841 | D429A | GAC-->GCC |
|  | Ck1527, CK732 | D429N | GAC-->AAC |
|  | CK1782 | V433D | GTC-->GAC |
|  | CK2194 | A424V | GCT-->GTT |

**Supplementary Table 2.**
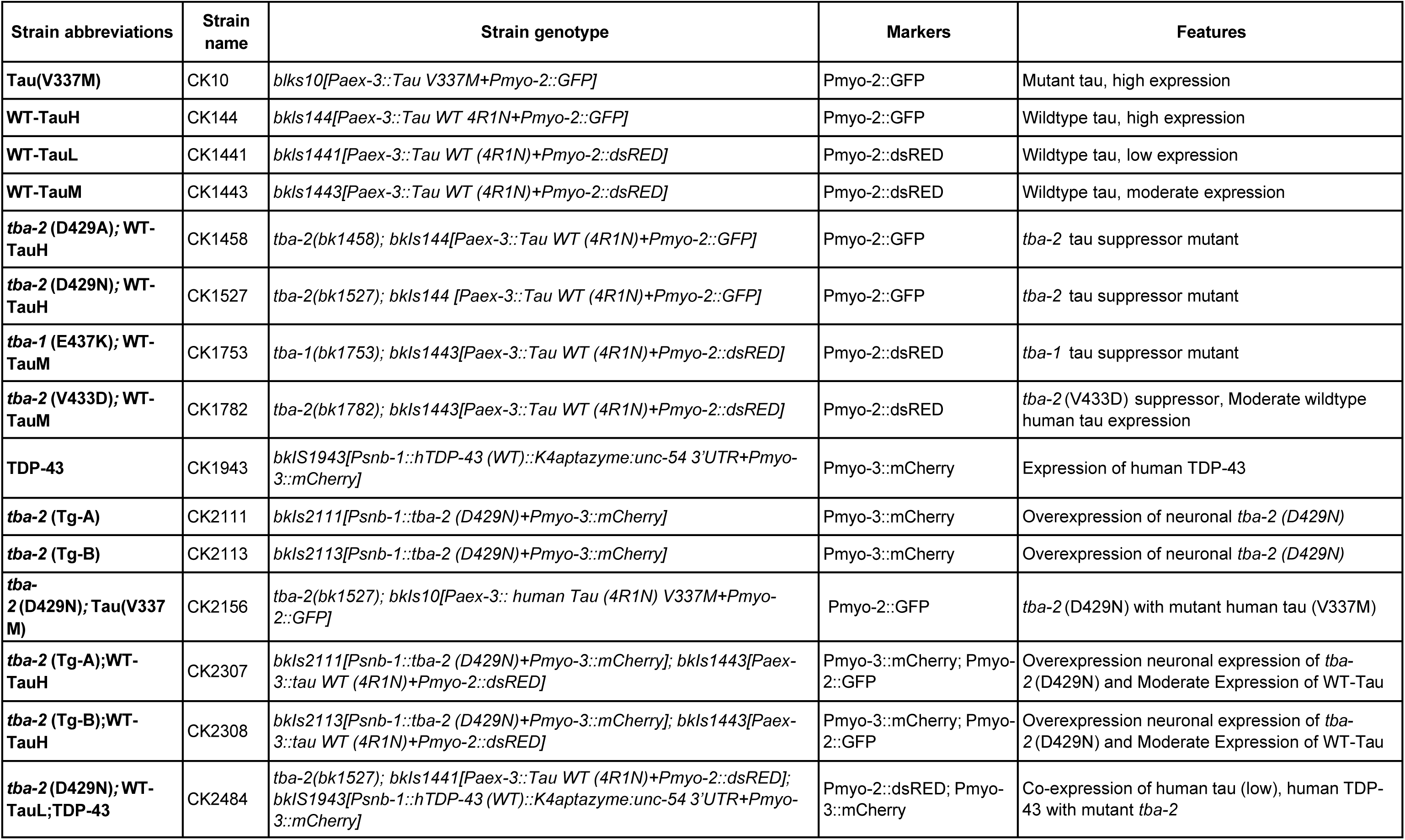
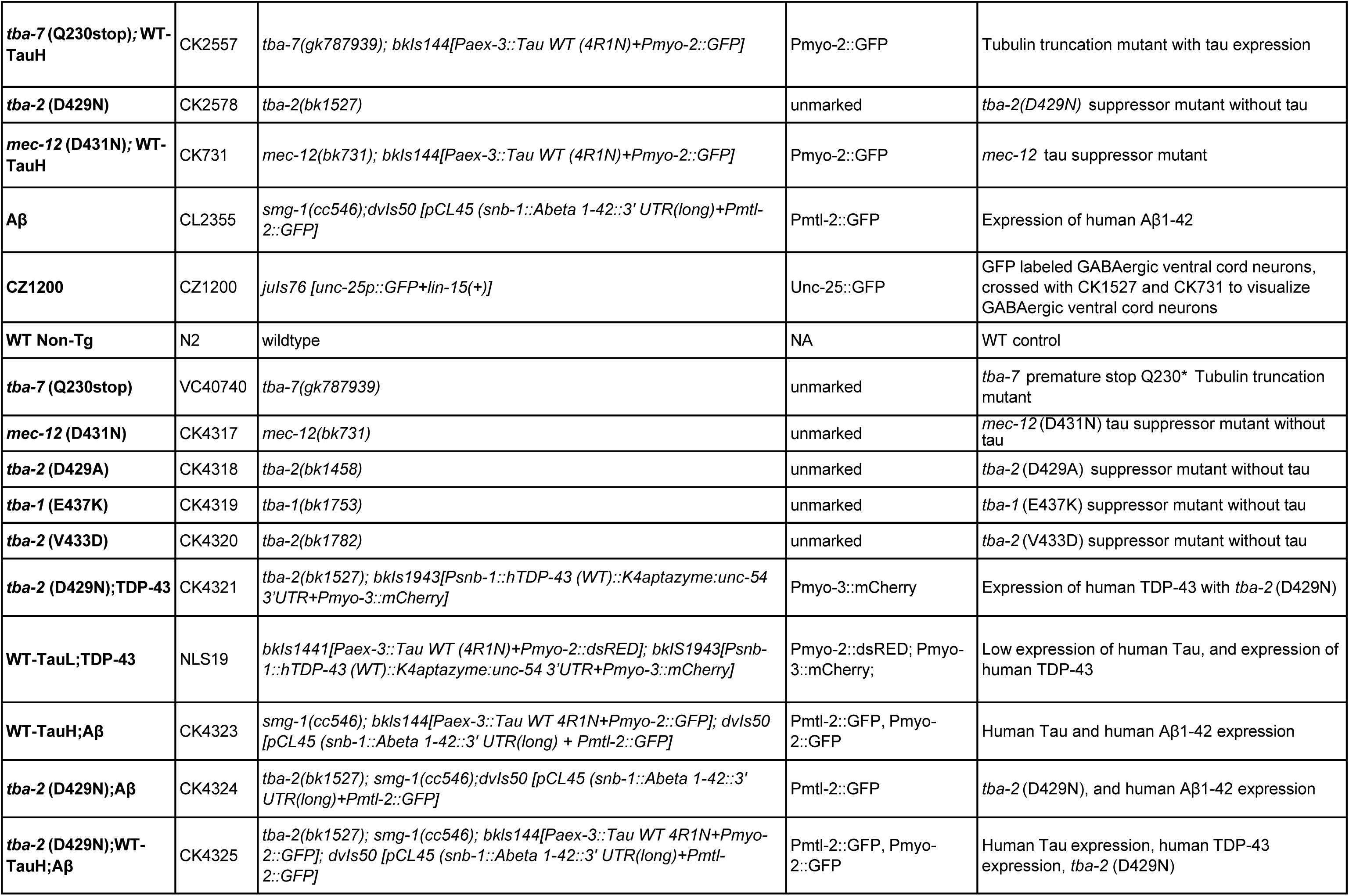
Strain List.

**Supplementary Table 3.** List of antibodies and dilutions used.

| Antibody | Antigen | Host Species | Source | Ref # | Application | Dilution |
| --- | --- | --- | --- | --- | --- | --- |
| Thermo SP70 | Pan-tau | Rabbit | ThermoFisher | MA5-16404 | Western Blot | Total Tau: 1:2500<br>RAB/RIPA: 1:2000<br>FA tau 1:1000 |
|  |  |  |  |  | PeggySue | 1:2000 |
| DAKO | Pan-tau | Rabbit | DAKO/Agilent | A0024(01-2) | Western blot | Total tau: 1:500,000<br>RAB/RIPA: 1:400,000<br>(Unsuitable for FA tau) |
|  |  |  |  |  | PeggySue | 1:10,000 |
| SP70 pAb | Pan-tau | Rabbit | Rockland | 200-c)1-b33 | Western Blot | Total Tau: 1:2500 |
| PHF1 | Phospho-tau p-ser396/Ser404 | Mouse | Peter Davies | NA | Western Blot | 1:1000 |
| CP13 | Phospho-tau | Mouse | Peter Davies | NA | Western Blot | 1:1000 |
| AT180 | Phospho-tau | Mouse | ThermoFisher | MN040 | Western Blot | 1:1000 |
| β-Tubulin | E7 mAb | Mouse | Developmental Studies Hybridoma Bank (Iowa City IA, USA) | NA | Western Blot | 1:5000 |
| 2° Ab mouse | Horseradish Peroxidase α-Ms IgG (H + L) | Goat | Jackson ImmunoResearch (West Grove, PA, USA) | 115-035-146 | Western Blot | 1:5000 |
|  |  |  |  |  | PeggySue | 1:100 |
| 2° Ab Rabbit | Horseradish Peroxidase α-Rb IgG (H + L) | Mouse | Jackson ImmunoResearch (West Grove, PA, USA) | 211-032-171 | Western Blot | 1:5000 |
|  |  |  |  |  | PeggySue | 1:100 |

**Supplementary Figure 1.**
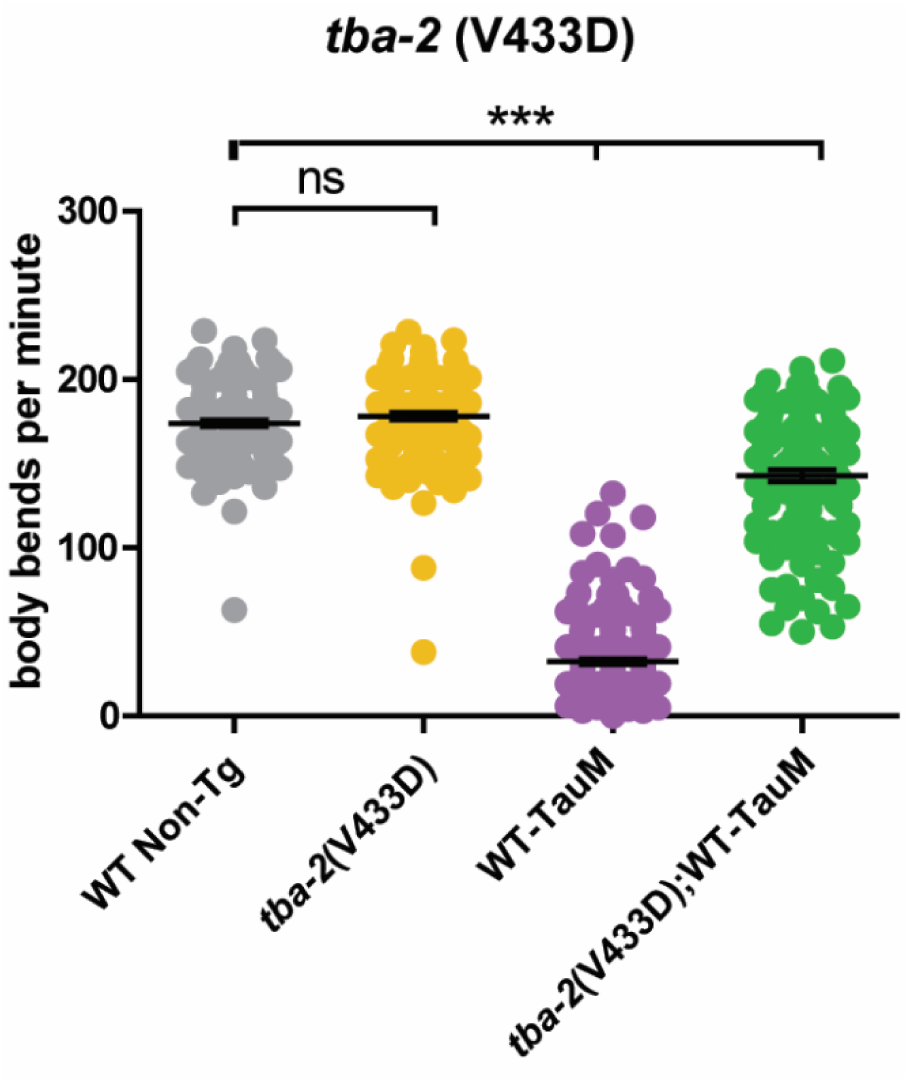
*tba-2*(V433D) shows moderate levels of suppression in a tau-transgenic strain expressing medium levels of wildtype human 4R1N tau (CK1443). (***p<0.0001, Kruskal-Wallis ANOVA with Dunn’s comparison, n≥114, error bars represent SEM).

**Supplementary Figure 2.**
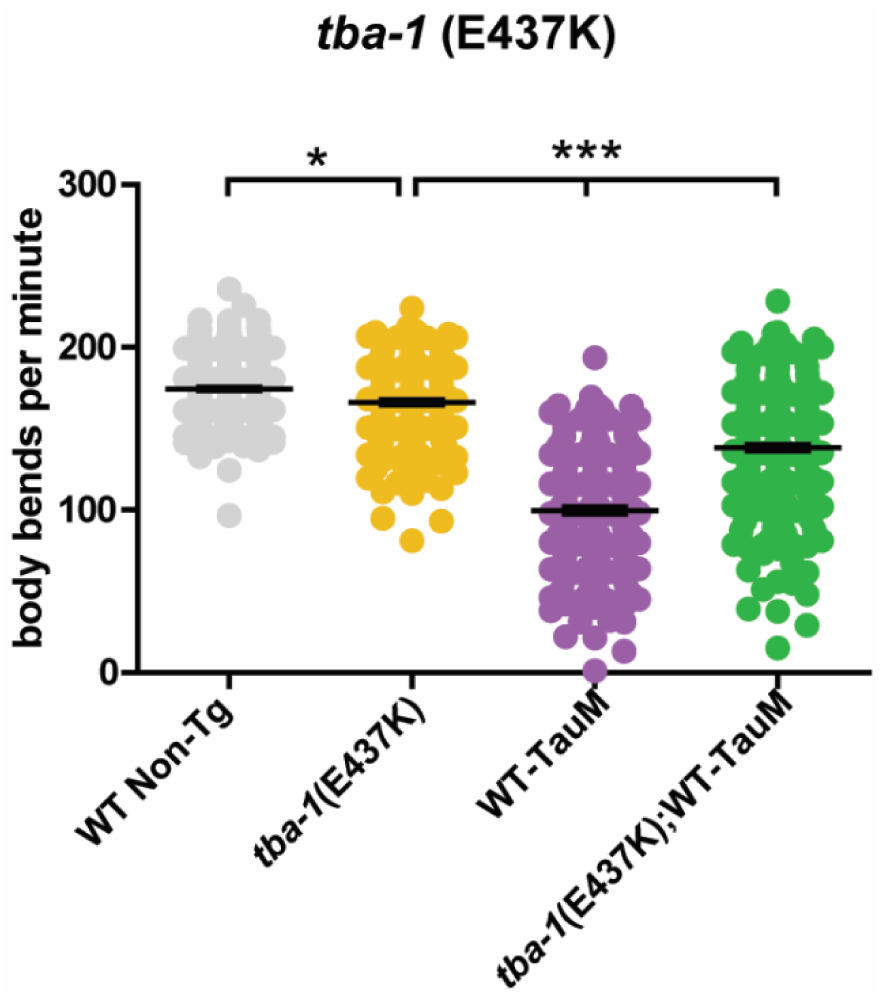
Mutant *tba-1* moderately suppresses tau-induced motility deficits. *tba-1*(E437K) shows moderate levels of suppression in a tau-transgenic strain expressing medium levels of wildtype human 4R1N tau (CK1443). (***p<0.0001, Kruskal-Wallis ANOVA with Dunn’s comparison, n≥230, error bars represent SEM).

**Supplementary Figure 3.**
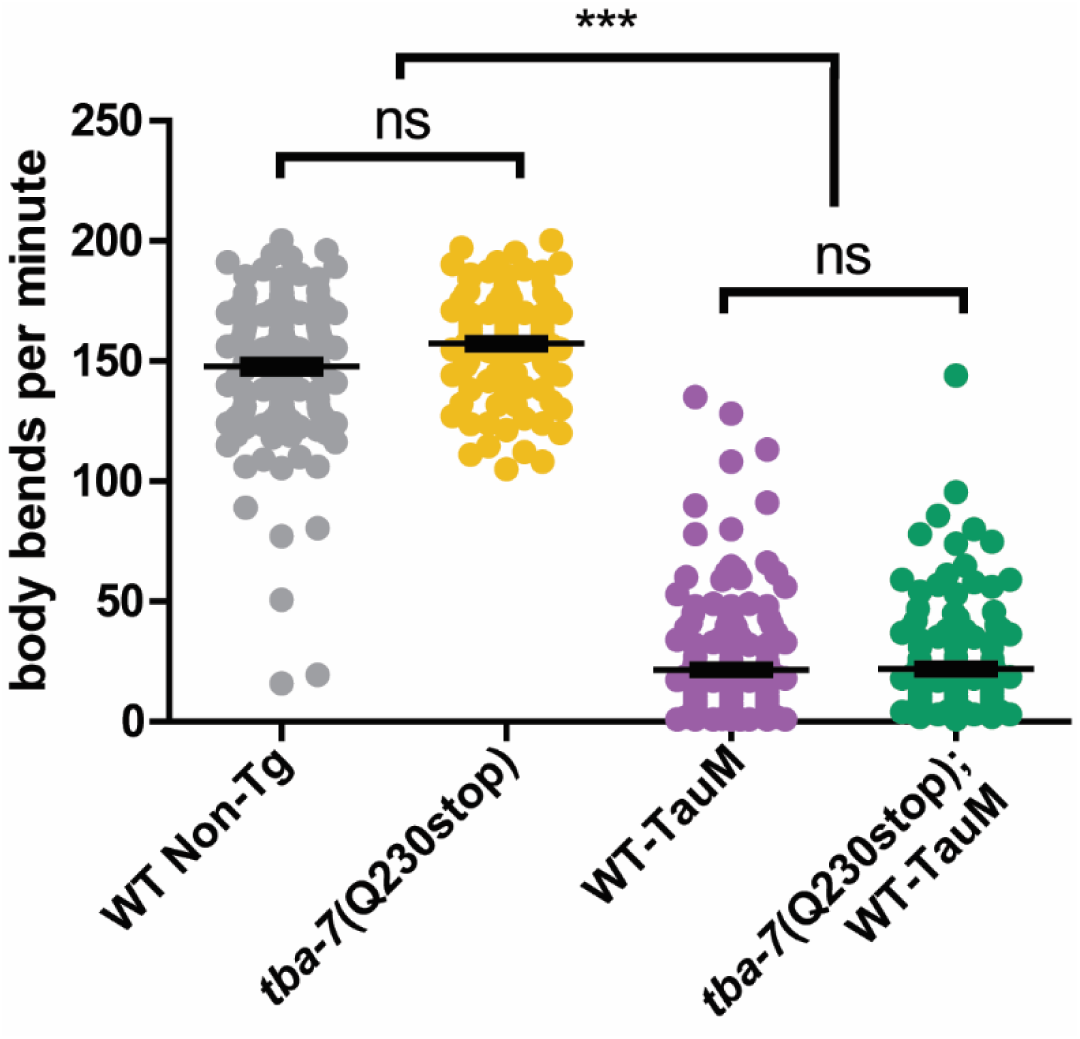
Truncated tubulin does not rescue tau-induced phenotypes. The tubulin truncation of *tba-7* at Q230 does not rescue tau-induced motility deficits in worms expressing wildtype human tau at moderate expression levels (CK1443). Kruskal-Wallis ANOVA with Dunn’s post-hoc test, \*\*\**p<0.0001*, n ≥ 146, error bars reflect SEM.

**Supplementary Figure 4.**
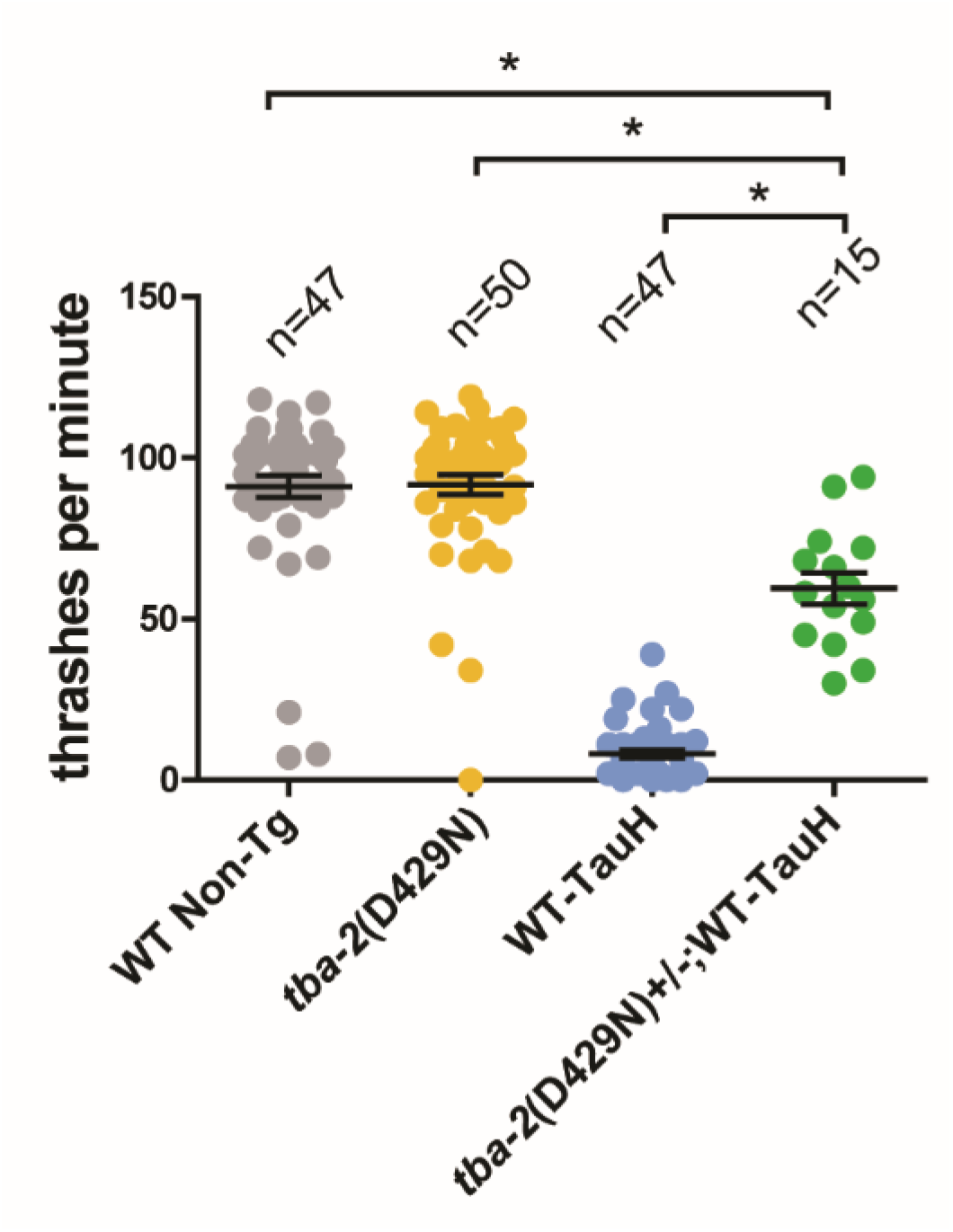
Tubulin mutations are strong semi-dominant suppressors of tau toxicity. *C. elegans* homozygous for tau, but heterozygous for the *tba-2(D429N)* were tested for homozygous tau suppression in a swimming assay. Population is low due to post-swim confirmation of heterozygosity. n ≥ 15, Kruskall-Wallis ANOVA with Dunn’s post-hoc test, *\*p<0.05* error bars reflect SEM.

**Supplementary Figure 5.**
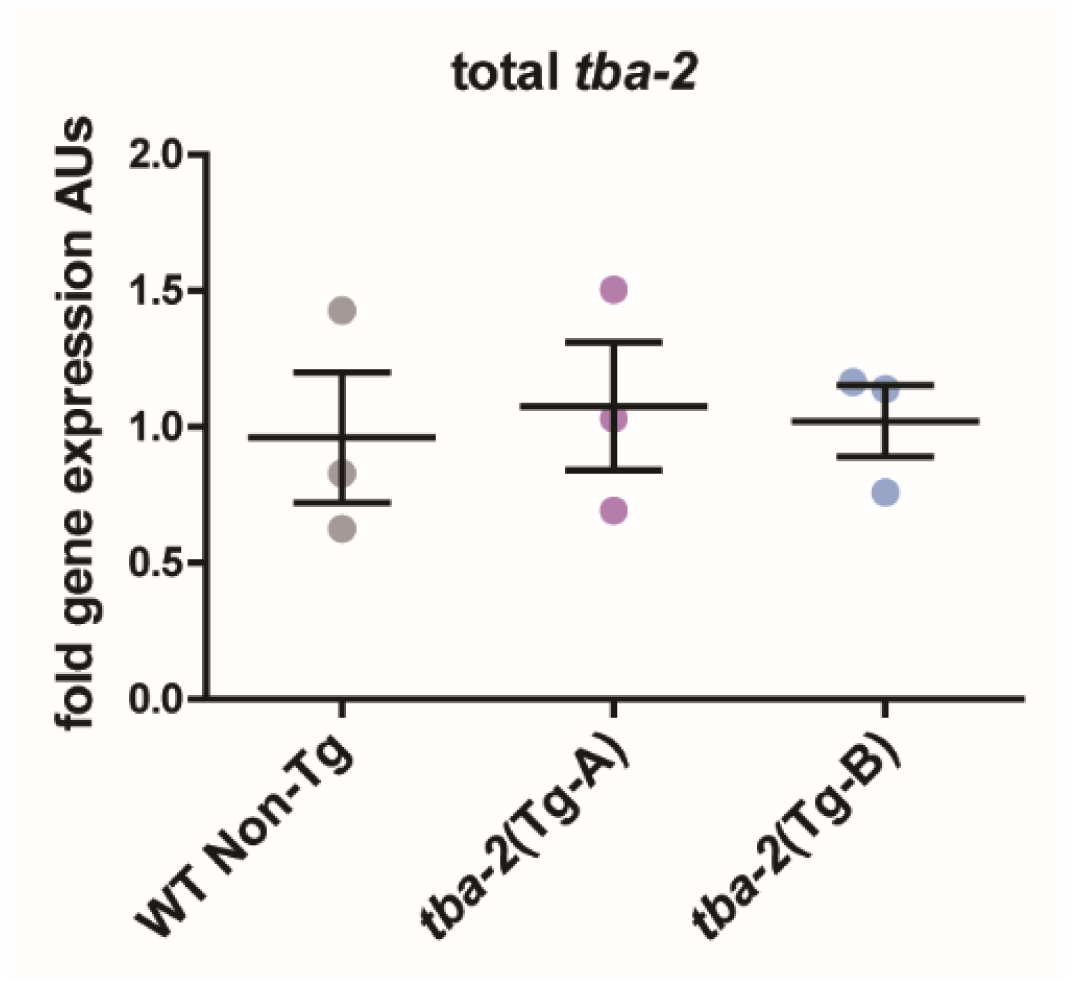
Pan-tissue control for total *tba-2* mRNA in qPCR experiments. Primers amplified endogenous *tba-2* expressed across all worm tissues which remains unchanged since overexpression of mutant tubulin is specific to neurons that make up only a small portion of total *C. elegans* cells.

**Supplementary Figure 6.**
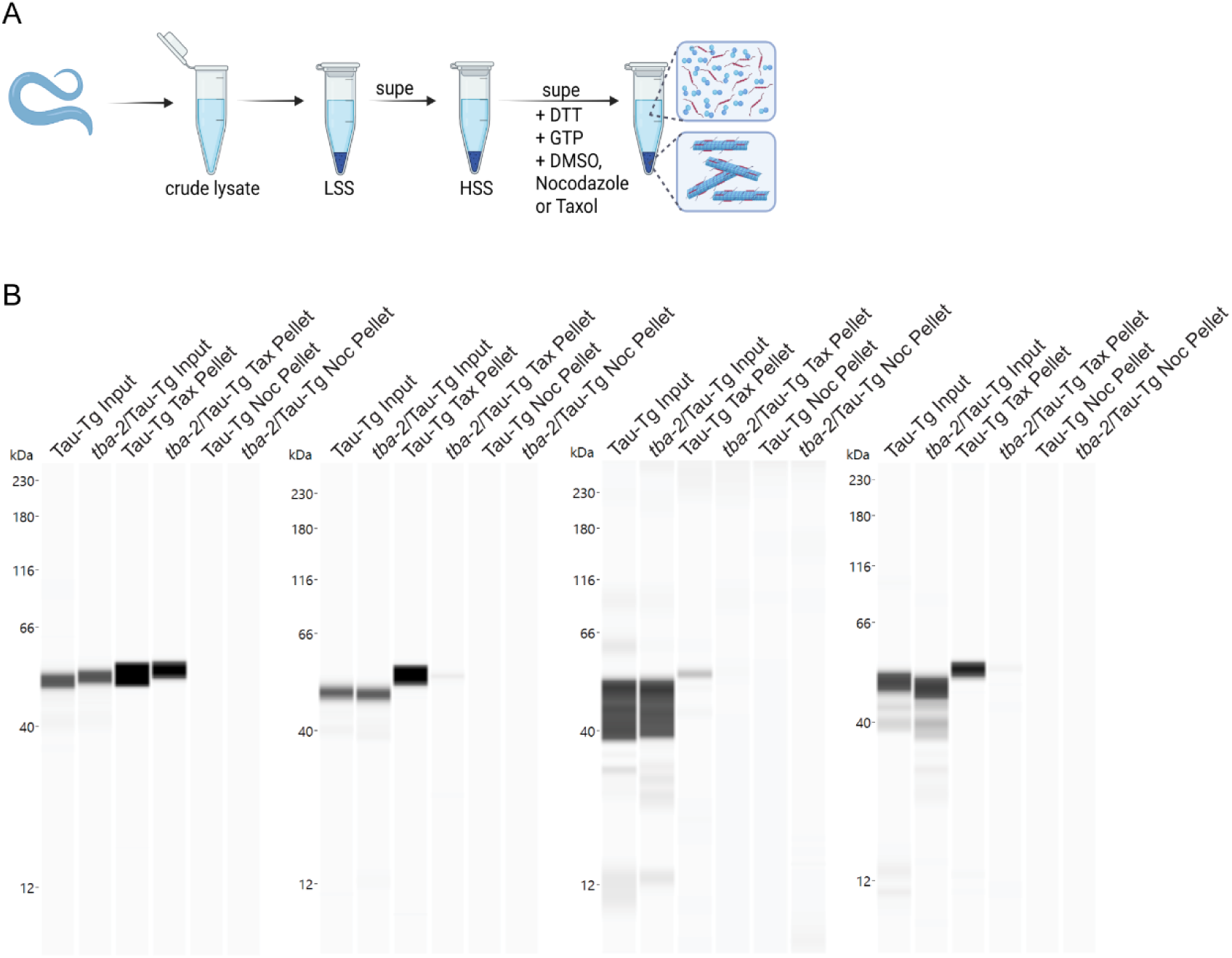
A. Workflow schematic for sedimentation assay [Created in BioRender. Wheeler, J. (2025) https://BioRender.com/fqyruiy] Biological replicates of capillary westerns data showing input tubulin, and sedimented microtubule mass after incubation with taxol or nocodazole.

**Supplementary Figure 7.**
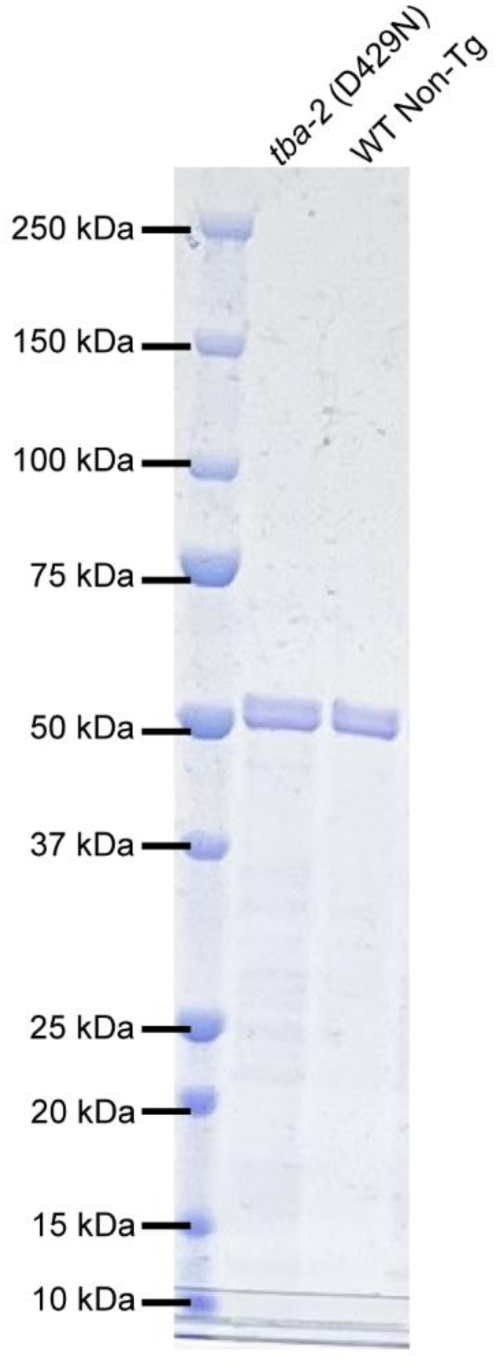
Representative Coomassie stained acrylamide gel showing approximately 0.2mg/ml tubulin isolated from *tba-2* (D429N) and WT Non-Tg animals.

## References

1. Götz J, Halliday G, Nisbet RM: Molecular Pathogenesis of the Tauopathies. Annu Rev Pathol 2019, 14:239–261.

2. Kneynsberg A, Combs B, Christensen K, Morfini G, Kanaan NM: Axonal Degeneration in Tauopathies: Disease Relevance and Underlying Mechanisms. Front Neurosci 2017, 11:572.

3. Dujardin S, Colin M, Buée L: Invited review: Animal models of tauopathies and their implications for research/translation into the clinic. Neuropathol Appl Neurobiol 2015, 41:59–80.

4. Spillantini MG, Murrell JR, Goedert M, Farlow MR, Klug A, Ghetti B: Mutation in the tau gene in familial multiple system tauopathy with presenile dementia. Proc Natl Acad Sci U S A 1998, 95:7737–7741.

5. Clark LN, Poorkaj P, Wszolek Z, Geschwind DH, Nasreddine ZS, Miller B, Li D, Payami H, Awert F, Markopoulou K, et al: Pathogenic implications of mutations in the tau gene in pallido-ponto-nigral degeneration and related neurodegenerative disorders linked to chromosome 17. Proc Natl Acad Sci U S A 1998, 95:13103–13107.

6. Hutton M, Lendon CL, Rizzu P, Baker M, Froelich S, Houlden H, Pickering-Brown S, Chakraverty S, Isaacs A, Grover A, et al: Association of missense and 5’-splice-site mutations in tau with the inherited dementia FTDP-17. Nature 1998, 393:702–705.

7. Dickson DW, Kouri N, Murray ME, Josephs KA: Neuropathology of frontotemporal lobar degeneration-tau (FTLD-tau). J Mol Neurosci 2011, 45:384–389.

8. Fitzpatrick AWP, Falcon B, He S, Murzin AG, Murshudov G, Garringer HJ, Crowther RA, Ghetti B, Goedert M, Scheres SHW: Cryo-EM structures of tau filaments from Alzheimer’s disease. Nature 2017, 547:185–190.

9. Goedert M: Cryo-EM structures of τ filaments from human brain. Essays Biochem 2021, 65:949–959.

10. Nilson AN, English KC, Gerson JE, Barton Whittle T, Nicolas Crain C, Xue J, Sengupta U, Castillo-Carranza DL, Zhang W, Gupta P, Kayed R: Tau Oligomers Associate with Inflammation in the Brain and Retina of Tauopathy Mice and in Neurodegenerative Diseases. J Alzheimers Dis 2017, 55:1083–1099.

11. Sferra A, Nicita F, Bertini E: Microtubule Dysfunction: A Common Feature of Neurodegenerative Diseases. Int J Mol Sci 2020, 21.

12. Alonso AD, Grundke-Iqbal I, Barra HS, Iqbal K: Abnormal phosphorylation of tau and the mechanism of Alzheimer neurofibrillary degeneration: sequestration of microtubule-associated proteins 1 and 2 and the disassembly of microtubules by the abnormal tau. Proc Natl Acad Sci U S A 1997, 94:298–303.

13. Brandt R, Bakota L: Microtubule dynamics and the neurodegenerative triad of Alzheimer&#x27;s disease: The hidden connection. Journal of Neurochemistry 2017, 143:409–417.

14. Hurd DD: Tubulins in C. elegans. In Wormbook. Edited by Community TCeR; 2018

15. Gasic I: Regulation of Tubulin Gene Expression: From Isotype Identity to Functional Specialization. Front Cell Dev Biol 2022, 10:898076.

16. Kadavath H, Hofele RV, Biernat J, Kumar S, Tepper K, Urlaub H, Mandelkow E, Zweckstetter M: Tau stabilizes microtubules by binding at the interface between tubulin heterodimers. Proc Natl Acad Sci U S A 2015, 112:7501–7506.

17. Best RL, LaPointe NE, Liang J, Ruan K, Shade MF, Wilson L, Feinstein SC: Tau isoform-specific stabilization of intermediate states during microtubule assembly and disassembly. J Biol Chem 2019, 294:12265–12280.

18. Breuzard G, Hubert P, Nouar R, De Bessa T, Devred F, Barbier P, Sturgis JN, Peyrot V: Molecular mechanisms of Tau binding to microtubules and its role in microtubule dynamics in live cells. Journal of cell science 2013, 126:2810–2819.

19. Barbier P, Zejneli O, Martinho M, Lasorsa A, Belle V, Smet-Nocca C, Tsvetkov PO, Devred F, Landrieu I: Role of Tau as a Microtubule-Associated Protein: Structural and Functional Aspects. Front Aging Neurosci 2019, 11:204.

20. Bunker JM, Wilson L, Jordan MA, Feinstein SC: Modulation of microtubule dynamics by tau in living cells: implications for development and neurodegeneration. Mol Biol Cell 2004, 15:2720–2728.

21. Goode BL, Denis PE, Panda D, Radeke MJ, Miller HP, Wilson L, Feinstein SC: Functional interactions between the proline-rich and repeat regions of tau enhance microtubule binding and assembly. Mol Biol Cell 1997, 8:353–365.

22. Hanger DP, Anderton BH, Noble W: Tau phosphorylation: the therapeutic challenge for neurodegenerative disease. Trends Mol Med 2009, 15:112–119.

23. Bramblett GT, Goedert M, Jakes R, Merrick SE, Trojanowski JQ, Lee VM: Abnormal tau phosphorylation at Ser396 in Alzheimer’s disease recapitulates development and contributes to reduced microtubule binding. Neuron 1993, 10:1089–1099.

24. Gustke N, Steiner B, Mandelkow EM, Biernat J, Meyer HE, Goedert M, Mandelkow E: The Alzheimer-like phosphorylation of tau protein reduces microtubule binding and involves Ser-Pro and Thr-Pro motifs. FEBS Lett 1992, 307:199–205.

25. Mandelkow EM, Schweers O, Drewes G, Biernat J, Gustke N, Trinczek B, Mandelkow E: Structure, microtubule interactions, and phosphorylation of tau protein. Ann N Y Acad Sci 1996, 777:96–106.

26. Zhang B, Carroll J, Trojanowski JQ, Yao Y, Iba M, Potuzak JS, Hogan AM, Xie SX, Ballatore C, Smith AB, 3rd, et al: The microtubule-stabilizing agent, epothilone D, reduces axonal dysfunction, neurotoxicity, cognitive deficits, and Alzheimer-like pathology in an interventional study with aged tau transgenic mice. J Neurosci 2012, 32:3601–3611.

27. Brunden KR, Zhang B, Carroll J, Yao Y, Potuzak JS, Hogan AM, Iba M, James MJ, Xie SX, Ballatore C, et al: Epothilone D improves microtubule density, axonal integrity, and cognition in a transgenic mouse model of tauopathy. J Neurosci 2010, 30:13861–13866.

28. Zhao X, Zeng W, Xu H, Sun Z, Hu Y, Peng B, McBride JD, Duan J, Deng J, Zhang B, et al: A microtubule stabilizer ameliorates protein pathogenesis and neurodegeneration in mouse models of repetitive traumatic brain injury. Sci Transl Med 2023, 15:eabo6889.

29. Kraemer BC, Zhang B, Leverenz JB, Thomas JH, Trojanowski JQ, Schellenberg GD: Neurodegeneration and defective neurotransmission in a <em>Caenorhabditis elegans</em> model of tauopathy. Proceedings of the National Academy of Sciences 2003, 100:9980–9985.

30. Guthrie CR, Greenup L, Leverenz JB, Kraemer BC: MSUT2 is a determinant of susceptibility to tau neurotoxicity. Hum Mol Genet 2011, 20:1989–1999.

31. Guthrie CR, Schellenberg GD, Kraemer BC: SUT-2 potentiates tau-induced neurotoxicity in Caenorhabditis elegans. Hum Mol Genet 2009, 18:1825–1838.

32. Kow RL, Sikkema C, Wheeler JM, Wilkinson CW, Kraemer BC: DOPA Decarboxylase Modulates Tau Toxicity. Biol Psychiatry 2018, 83:438–446.

33. Eck RJ, Kow RL, Black AH, Liachko NF, Kraemer BC: SPOP loss of function protects against tauopathy. Proceedings of the National Academy of Sciences 2023, 120:e2207250120.

34. Kow RL, Black AH, Henderson BP, Kraemer BC: Sut-6/NIPP1 modulates tau toxicity. Hum Mol Genet 2023, 32:2292–2306.

35. Kraemer BC, Schellenberg GD: SUT-1 enables tau-induced neurotoxicity in C. elegans. Hum Mol Genet 2007, 16:1959–1971.

36. LeBlanc Katherine R, Eck Randall J, Saxton Aleen D, McMillan Pamela J, Wheeler Jeanna M, Liachko Nicole F, Keene C D, Latimer Caitlin S, Kow Rebecca L, Kraemer Brian C: Tri-snRNP activity modulates tauopathy phenotypes. NAR Molecular Medicine 2025, 2.

37. Hoyle HD, Raff EC: Two Drosophila beta tubulin isoforms are not functionally equivalent. J Cell Biol 1990, 111:1009–1026.

38. Hutchens JA, Hoyle HD, Turner FR, Raff EC: Structurally similar Drosophila alpha-tubulins are functionally distinct in vivo. Mol Biol Cell 1997, 8:481–500.

39. Panda D, Miller HP, Banerjee A, Ludueña RF, Wilson L: Microtubule dynamics in vitro are regulated by the tubulin isotype composition. Proc Natl Acad Sci U S A 1994, 91:11358–11362.

40. Pucciarelli S, Ballarini P, Sparvoli D, Barchetta S, Yu T, Detrich HW, 3rd, Miceli C: Distinct functional roles of β-tubulin isotypes in microtubule arrays of Tetrahymena thermophila, a model single-celled organism. PLoS One 2012, 7:e39694.

41. Hammarlund M, Hobert O, Miller DM, 3rd, Sestan N: The CeNGEN Project: The Complete Gene Expression Map of an Entire Nervous System. Neuron 2018, 99:430–433.

42. Benbow SJ, Strovas TJ, Darvas M, Saxton A, Kraemer BC: Synergistic toxicity between tau and amyloid drives neuronal dysfunction and neurodegeneration in transgenic C. elegans. Hum Mol Genet 2020, 29:495–505.

43. Latimer CS, Stair JG, Hincks JC, Currey HN, Bird TD, Keene CD, Kraemer BC, Liachko NF: TDP-43 promotes tau accumulation and selective neurotoxicity in bigenic Caenorhabditis elegans. Dis Model Mech 2022, 15.

44. Kow RL, Black AH, Saxton AD, Liachko NF, Kraemer BC: Loss of aly/ALYREF suppresses toxicity in both tau and TDP-43 models of neurodegeneration. Geroscience 2022, 44:747–761.

45. Wang S, Wu D, Quintin S, Green RA, Cheerambathur DK, Ochoa SD, Desai A, Oegema K: NOCA-1 functions with γ-tubulin and in parallel to Patronin to assemble non-centrosomal microtubule arrays in C. elegans. Elife 2015, 4:e08649.

46. Messina S, Sframeli M: New Treatments in Spinal Muscular Atrophy: Positive Results and New Challenges. J Clin Med 2020, 9.

47. Chilcott EM, Muiruri EW, Hirst TC, Yáñez-Muñoz RJ: Systematic review and meta-analysis determining the benefits of in vivo genetic therapy in spinal muscular atrophy rodent models. Gene Therapy 2022, 29:498–512.

48. Brenner S: The genetics of Caenorhabditis elegans. Genetics 1974, 77:71–94.

49. De Stasio EA, Dorman S: Optimization of ENU mutagenesis of Caenorhabditis elegans. Mutat Res 2001, 495:81–88.

50. Waldherr SM, Strovas TJ, Vadset TA, Liachko NF, Kraemer BC: Constitutive XBP-1s-mediated activation of the endoplasmic reticulum unfolded protein response protects against pathological tau. Nature Communications 2019, 10:4443.

51. Portman DS: Profiling C. elegans gene expression with DNA microarrays. In WormBook. Edited by Community TCeR: WormBook; 2006

52. Schindelin J, Arganda-Carreras I, Frise E, Kaynig V, Longair M, Pietzsch T, Preibisch S, Rueden C, Saalfeld S, Schmid B: Fiji: an open-source platform for biological-image analysis. Nature methods 2012, 9:676–682.

53. McDermott JB, Aamodt S, Aamodt E: ptl-1, a Caenorhabditis elegans gene whose products are homologous to the tau microtubule-associated proteins. Biochemistry 1996, 35:9415–9423.

54. Widlund PO, Podolski M, Reber S, Alper J, Storch M, Hyman AA, Howard J, Drechsel DN: One-step purification of assembly-competent tubulin from diverse eukaryotic sources. Mol Biol Cell 2012, 23:4393–4401.

55. McMillan PJ, Benbow SJ, Uhrich R, Saxton A, Baum M, Strovas T, Wheeler JM, Baker J, Liachko NF, Keene CD, et al: Tau-RNA complexes inhibit microtubule polymerization and drive disease-relevant conformation change. Brain 2023, 146:3206–3220.

56. Baker JD, Uhrich RL, Kraemer GC, Love JE, Kraemer BC: A drug repurposing screen identifies hepatitis C antivirals as inhibitors of the SARS-CoV2 main protease. PLoS One 2021, 16:e0245962.

